# Quantitative Comparison of 3D-1D Vascular Coupling Models: Lateral Average versus Sphere of Influence Methods

**DOI:** 10.64898/2026.07.12.738114

**Authors:** Rohan Amare, Duygu Vargun, Pierce Zhang, Steve Parrish, Danielle Stolley, Chimamanda Santos, Megan Jacobsen, Erik Cressman, Beatrice Riviere, David T. Fuentes

## Abstract

Computational models coupling one-dimensional vascular networks with three-dimensional tissue domains are widely used for predicting blood flow distribution in tumor perfusion, drug delivery, and therapeutic planning. Two prominent coupling paradigms have emerged: the Lateral Average Model (LAM) which implements distributed transmural exchange via a vessel wall conductivity parameter *γ* (m Pa^−1^ s^−1^), and the Sphere of Influence (SOI) model, which employs localized terminal coupling via a source sphere radius *ε* (m). Despite their broad application, systematic quantitative comparisons of their parametric behavior and predictive equivalence remain lacking.

We compare LAM and SOI in 3D-1D simulations on a benchmark vascular network and a porcine liver study with a hepatic arterial network reconstructed from CT arteriography.

Across a benchmark vascular network under three sink configurations, the LAM net flow rate rose smoothly with *γ* and saturated at a plateau, while the SOI net flow rate increased with *ε* without saturating; as a result, global-flow equivalence between the two formulations exists only for particular boundary geometries, and not at all within the tested parameter range for one of the three configurations examined. Despite this partial agreement in total flow, the two models diverged substantially in regional perfusion: in a porcine hepatic arterial network reconstructed from CT arteriography, SOI predicted stable perfusion fractions to two regions of interest across its full tested parameter range, whereas LAM predictions for the same regions varied several-fold with vessel wall permeability and, at low permeability, could invert which region received more flow. These results indicate that the choice of coupling model has limited consequence for predicted total organ flow but substantial consequence for predicted local drug delivery, and we provide guidance for selecting between the two formulations depending on the clinical or research question being asked.

**Author Summary:** When doctors plan treatments for liver cancer, they often rely on computer simulations to predict how blood flows through the liver and how well a drug will reach the tumor. These simulations depend on mathematical models that describe how blood moves from vessels into surrounding tissue. Two commonly used approaches exist for building these models, but researchers have generally chosen between them based on habit or convenience rather than on a principled understanding of how their predictions differ.

In this work, we directly compared these two approaches, one that spreads blood exchange continuously along the vessel wall, and one that delivers blood from the vessel tips into a surrounding spherical zone, using both a simple test network and a realistic pig liver reconstructed from medical imaging. We found that the two approaches can agree on the total amount of blood reaching the liver, but disagree substantially on *where* that blood goes within the tissue. This distinction matters enormously for treatment planning: a model that predicts the right total blood flow but delivers it to the wrong region of the liver could lead to an inaccurate forecast of drug concentration at the tumor site. Our results provide practical guidance for researchers on which approach to use depending on what information is available and what question is being asked.

## Introduction

Computational modeling of blood flow in biological systems has emerged as an indispensable tool for understanding complex hemodynamic phenomena ranging from large-scale cardiovascular function to microscopic tissue perfusion [**?, ?**]. The inherent multiscale nature of vascular networks—spanning from centimeter-scale arteries to micrometer-scale capillaries—poses fundamental modeling challenges that have motivated the development of geometrical multiscale approaches coupling models of different dimensional complexity [**?, ?**]. Among these approaches, the coupling of three-dimensional (3D) tissue domains with embedded one-dimensional (1D) vascular networks has proven particularly valuable for applications where detailed tissue-scale phenomena must be resolved while maintaining computational tractability for large vascular trees [**?, ?**]. Such 3D-1D coupled models have found widespread application in tumor perfusion analysis [**?, ?**], drug delivery optimization [**?, ?**], tissue engineering [**?**], and therapeutic planning for interventional procedures [**?**].

The conceptual foundation of 3D-1D coupling rests on representing blood vessels as one-dimensional concentrated sources or sinks embedded within a three-dimensional continuum governed by Darcy’s law or Stokes equations [**?**]. This geometrical reduction decreases computational cost by eliminating the need to explicitly mesh small vessel lumens while retaining essential coupling between vascular and tissue compartments [**?, ?**]. However, the implementation of this coupling involves critical modeling choices regarding how mass, momentum, and energy are exchanged between dimensional domains. Two prominent paradigms have emerged: the Lateral Average Model (LAM) and the Sphere of Influence (SOI) model. Both approaches utilize the Hagen-Poiseuille equation in the 1D vessels and the Darcy’s law for flow through porous media in the 3D tissue domain. However, LAM and SOI are fundamentally different with respect to the coupling mechanisms between the 1D vessels and the 3D tissue.

The Lateral Average Model implements distributed transmural exchange along the entire length of each vessel segment [**?, ?**]. In this formulation, the coupling between domains is achieved through a vessel wall permeability parameter *γ* that characterizes the resistance to lateral fluid exchange across the vessel wall. Mathematically, this coupling appears as a distributed line-source term in the 3D tissue equations and a corresponding sink term in the 1D vessel equations, with magnitudes proportional to the vessel wall permeability and the pressure difference between vessels and tissue [**?, ?**]. Additional coupling occurs at the terminal nodes of the vessels, mathematically characterized as point sources in the 3D tissue and Neumann boundary conditions in the 1D vessels. The LAM has been successfully applied to model plasma filtration in arterial walls [**?**], drug eluting stent performance [**?**], and microvascular exchange in complex tissue architectures [**?**]. Its distributed coupling mechanism naturally captures transmural exchange processes that occur continuously along vessel lengths, making it particularly appropriate for modeling scenarios where vessel wall permeability plays a dominant physiological role, such as inflammation-induced hyperpermeability, tumor-associated vascular leakage, or pharmacologically-induced vascular normalization [**?**].

In contrast, the Sphere of Influence model employs localized coupling exclusively at vessel terminals through Dirac delta functions [**?, ?**]. In this approach, blood delivered by the 1D vascular network is transferred to the 3D tissue domain only at terminal nodes, with the exchange distributed over a spherical region of radius *ε* centered at each terminal node. The coupling is mathematically represented using a Dirac distribution function *η*_*ε*_(*x* − *y*) that weights the contribution of each terminal to nearby tissue voxels based on distance, with the sphere of influence radius *ε* controlling the spatial extent of the exchange zone [**?**]. This formulation is motivated by the physiological observation that in many vascular beds, particularly those dominated by tree-like arterial networks, the primary site of blood-tissue exchange occurs at the capillary level where arterioles terminate and distribute blood into dense capillary plexuses [**?**]. The SOI model has been successfully employed in whole-organ perfusion simulations [**?**], multiscale thermoregulation modeling [**?**], and patient-specific coronary hemodynamics [**?**]. Its computational efficiency stems from eliminating the need to track distributed exchange along potentially millions of vessel segments, instead concentrating coupling computations at a smaller number of terminal locations.

Despite the widespread adoption of both LAM and SOI approaches, a systematic quantitative comparison of these coupling paradigms remains absent from the literature. Existing studies typically employ one model or the other based on historical precedent, computational convenience, or qualitative physical intuition, without rigorous investigation of how model choice affects predicted hemodynamics [**?**]. Several fundamental questions remain unresolved: Can LAM and SOI produce equivalent hemodynamic predictions, and if so, what relationships must exist between their respective coupling parameters (*γ* and *ε*)? How sensitive are flow predictions to parameter variations in each model, and what implications does this sensitivity have for parameter estimation from limited clinical data? Do the models exhibit different spatial redistribution of blood flow in response to parameter changes, and how might this affect predictions for heterogeneous pathological vasculature? Which model is more appropriate for specific physiological scenarios or clinical applications?

The resolution of these questions has immediate practical significance for computational hemodynamics practitioners. In tumor perfusion modeling, where vascular heterogeneity and abnormal permeability are hallmarks of the pathological microenvironment [**?, ?**], the choice between distributed lateral exchange (LAM) and terminal coupling (SOI) could fundamentally alter predictions of drug delivery, hypoxia distribution, and therapeutic response [**?**]. In tissue engineering applications, where vascularization strategies must be optimized to ensure adequate perfusion of engineered constructs [**?, ?**], understanding the relationship between coupling mechanisms and regional flow patterns is essential for rational scaffold design. In patient-specific cardiovascular modeling, where parameter estimation from sparse clinical imaging data is unavoidable [**?**], knowledge of parameter sensitivity and uncertainty propagation characteristics becomes critical for establishing predictive reliability [**?**].

This study presents a comprehensive quantitative comparison of the Lateral Average Model and Sphere of Influence coupling approaches using finite element and finite volume discretizations applied to a physiologically-motivated vascular network embedded in tissue domains. We systematically investigate three key aspects of model behavior: (1) equivalence relationships between *γ* and *ε* that produce identical total domain flow rates, (2) sensitivity of flow predictions to parameter variations in each model, and (3) spatial redistribution of regional tissue perfusion in response to coupling parameter changes i.e. 3D pressure map and local flux variations. Our analysis employs a controlled computational framework where both models are implemented with identical discretization schemes, boundary conditions, and vessel network geometry, isolating the effects of coupling mechanism from confounding numerical or geometric factors.

## Materials and methods

**Fig 1.**
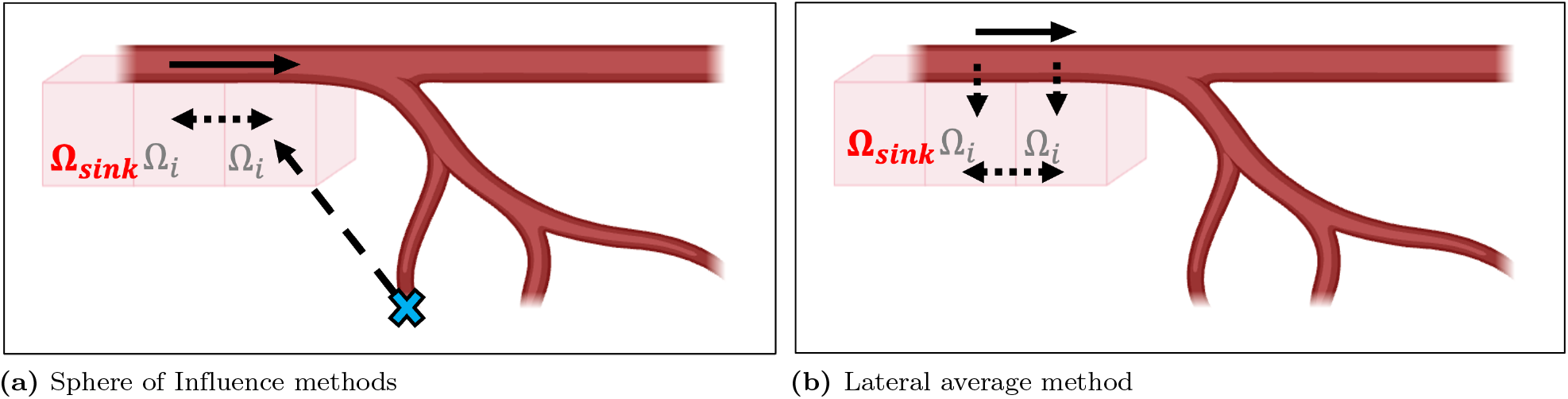
Illustration of the two mathematical coupling models. (a) Sphere of Influence Method. The coupling happens at the end of the segmented 1D blood vessels with the 3D porous tissue. The region Ω_*sink*_ represents the voxel where the sink boundary condition is applied. (b) Lateral average method. The coupling happens along the wall vessels using a lateral average operator and there is also coupling at the terminal nodes.

### Sphere of Influence Method

#### Physical Motivation

The Sphere of Influence (SOI) method addresses the fundamental challenge of coupling discrete one-dimensional vascular networks with three-dimensional tissue domains where explicit capillary geometry cannot be resolved. Rather than modeling the infinitely complex sub-resolution capillary architecture explicitly, the SOI method distributes flow from each terminal node over a finite spherical volume, effectively representing the collective behavior of unresolved branching vessels. The characteristic radius *ε* represents the spatial extent of unresolved capillary networks extending from each terminal node.

#### 3D Tissue Model Problem

The tissue is a 3D domain Ω partitioned into a set of voxels {Ω_*i*_}.

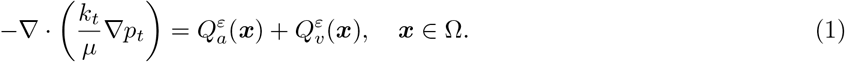

In the partial differential equation (Eq. (1)), the fluid viscosity is denoted by *µ >* 0 [Pa·s] and the permeability of the tissue is denoted by *k*_*t*_ *>* 0 [m^2^]. In (1), the volumetric source and sink functions are denoted by 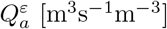 and 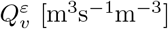 respectively, representing blood delivered from arterial terminals and collected by venous terminals.

Assume we have *N*_*out*_ terminal nodes for the arterial blood vessels and denote them by 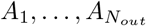. The spatial coordinates of node *A*_*k*_ are denoted by ***x***_*k*_ ∈ ℝ^3^. Let *B*_*ε*_(*A*_*k*_) = {***x*** ∈ Ω : ∥***x*** − ***x***_*k*_∥ *< ε*} be the sphere of radius *ε* and center *A*_*k*_.

#### Regularized Distribution Function

The total volumetric source at position ***x*** is given as:

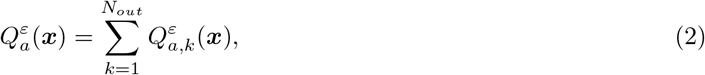

where *Q*^*ε*^ corresponds to the source term from a single arterial terminal node *A*_*k*_:

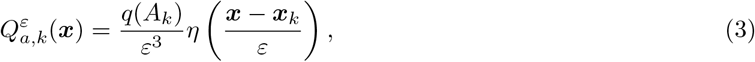

with *η*(***x***) a compactly supported, smooth distribution function defined below. The scaling factor 1*/ε*^3^ ensures dimensional consistency [m^−3^], transforming the dimensionless shape function *η* into a volumetric distribution.

The distribution function *η* : R^3^ → R is chosen as:

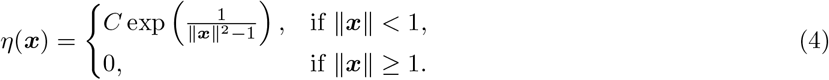

This exponential form provides several critical properties:

- **Compact support:** *η*(***x***) = 0 for ∥ ***x*** ∥ ≥ 1, ensuring flow distribution is strictly localized within the sphere of influence.
- **Smoothness:** *η* ∈ *C*^∞^(R^3^), eliminating spurious pressure oscillations and ensuring well-posed finite element formulations.
- **Physical realism:** Exponential decay provides stronger weighting to voxels closer to the terminal node.

#### Normalization and Mass Conservation

The constant *C* in (4) is chosen to satisfy the normalization condition:

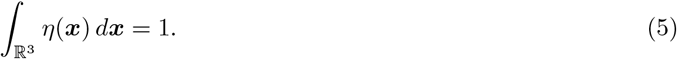

This integral is typically evaluated numerically to determine *C* for the three-dimensional case. The normalization ensures that the total mass delivered from terminal node *A*_*k*_ equals the flow rate *q*(*A*_*k*_):

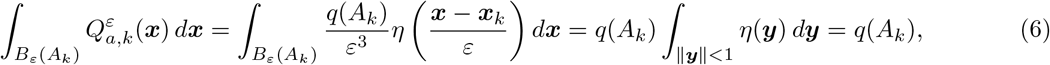

where we used the change of variables ***y*** = (***x*** *™* ***x***_*k*_)*/ε* and *d****x*** = *ε*^3^*d****y***.

This demonstrates **global mass conservation**: all flow exiting the terminal vessel node is distributed to the surrounding tissue.

We observe that

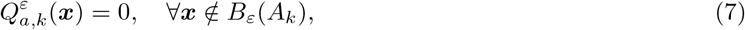

ensuring computational efficiency by limiting coupling to local regions.

#### Venous Sink Term

For the sink term 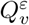, we define a specific drainage region in the domain and denote it by Ω_*sink*_ ⊂ Ω:

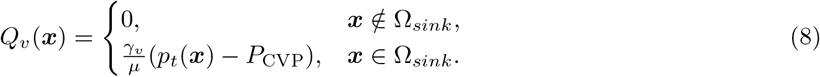

The central venous pressure *P*_CVP_ [Pa] is a prescribed positive constant representing the venous outflow boundary condition. The parameter *γ*_*v*_ [m^−1^] controls the drainage conductance from tissue to the venous system.

#### 3D Tissue Finite Volume Discretization

The domain Ω is partitioned into cubic voxels (with side length *ℓ* = 1 mm), denoted by Ω_*i*_ with volume |Ω_*i*_| = *ℓ*^3^. We integrate Eq. (1) over Ω_*i*_ and apply the divergence theorem:

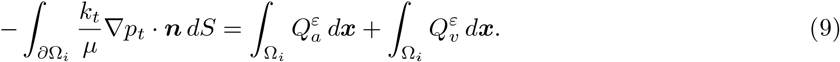

Next, we approximate the normal derivatives by centered finite differences. Let *p*_*t,i*_ denote the constant unknown approximating the pressure *p*_*t*_ at the center of voxel Ω_*i*_. For neighboring voxels Ω_*i*_ and Ω_*j*_ sharing a face *γ*_*ij*_:

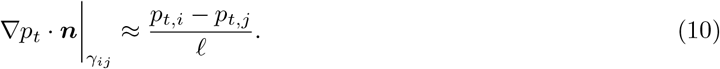

Let *I*_*i*_ denote the set of indices *j* such that Ω_*i*_ and Ω_*j*_ share a face (i.e., they are neighbors). For a cubic lattice with uniform spacing, |*I*_*i*_| = 6 for interior voxels. The discrete flux through all faces becomes:

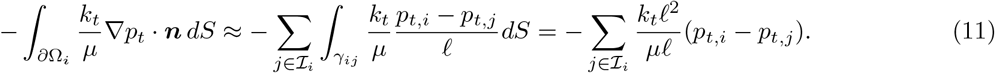

Note that each face has area *ℓ*^2^ and the finite difference uses spacing *ℓ*, yielding:

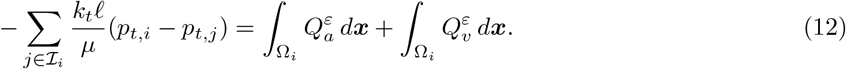

##### Venous sink discretization

Approximating *Q*_*v*_ as piecewise constant, we have

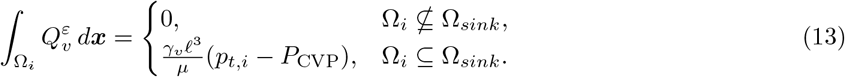

##### Arterial source discretization

Let ***x***_*i*_ denote the center of voxel Ω_*i*_. Using midpoint quadrature:

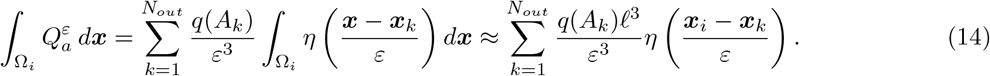

Define the normalized distribution weights:

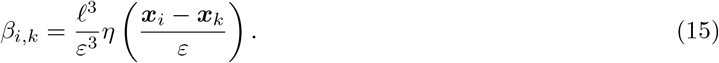

These weights satisfy **discrete mass conservation**:

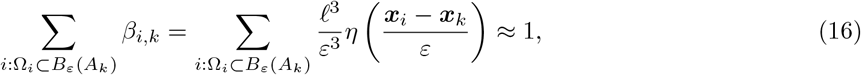

with quadrature error *O*(*ℓ*^2^) for smooth *η*.

##### Complete discrete tissue equations

If voxel Ω_*i*_ ⊆ Ω_*sink*_:

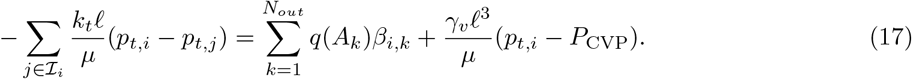

If voxel Ω_*i*_ ⊈ Ω_*sink*_:

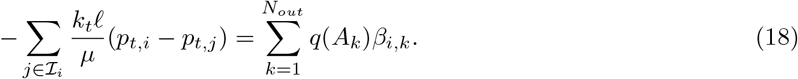

#### 1D Vasculature Discrete Problem

The vasculature is represented as a network graph G = (V, E) where V is the set of nodes (bifurcation points and terminals) and E is the set of edges (vessel segments). Each vessel segment is further partitioned into intervals and each interior node of the partition has an associated pressure *P*_*i*_ [Pa].

Using the Hagen-Poiseuille law for laminar flow in a cylindrical vessel, the volumetric flow rate *Q*_*ij*_ [m^3^/s] between nodes *P*_*i*_ and *P*_*j*_ connected by edge *e*_*ij*_ ∈ E is:

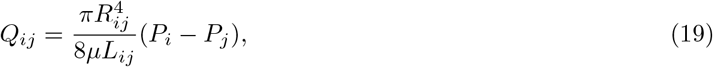

where *R*_*ij*_ [m] is the vessel radius and *L*_*ij*_ [m] is the vessel length. The coefficient 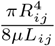 represents the hydraulic conductance [m^3^Pa^−1^s^−1^] of the vessel segment.

For an interior node *P*_*i*_ ∈ *V*_*I*_ (neither inlet nor terminal), let *N*_*i*_ denote the set of neighboring nodes directly connected to *P*_*i*_. Mass conservation (Kirchhoff’s current law) at node *P*_*i*_ requires:

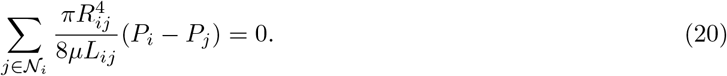

For an inlet node *P*_*i*_ ∈ *V*_*in*_, we prescribe the pressure as a Dirichlet boundary condition:

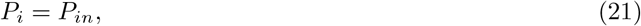

where *P*_*in*_ [Pa] is a specified constant (e.g., mean arterial pressure).

#### Coupling Between 3D and 1D Discrete Problems

For each arterial terminal node *A*_*k*_ ∈ *V*_*T*_, let *P*_*k*_ denote the vessel pressure at that terminal. The flow rate *q*(*A*_*k*_) exiting the terminal is coupled to the vessel pressure and the surrounding tissue pressure through a pressure-driven relationship. The coupling equation is:

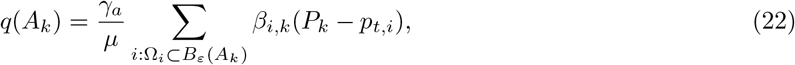

where *γ*_*a*_ [m Pa^−1^ s^−1^] is an effective conductance parameter representing the collective resistance of unresolved capillaries between the terminal and tissue.

##### Physical interpretation

Equation (22) states that flow from the terminal is driven by the pressure difference between the vessel and the weighted average tissue pressure within the sphere of influence. The weights *β*_*i,k*_ defined in Eq. (15) ensure that voxels closer to the terminal contribute more strongly to this average.

### Lateral Average Model

We now formulate a model that couples the centerlines of the blood vessels with the surrounding tissue: the model was introduced by [**?**] for different boundary conditions. The unknowns are the scalar fields, tissue pressure *p*_*t*_, and blood pressure *P*. The set of edges *E* in the network graph *G* of the vasculature consists of *N* edges *e*_*j*_. Each graph edge *e*_*j*_ of the network, namely the line segment between two bifurcation points, is parametrized by a function *λ*_*j*_(*s*) and it corresponds to the centerline of a blood vessel with cross section Θ_*j*_(*s*) (and area |Θ_*j*_(*s*)| and perimeter |∂Θ_*j*_(*s*)|).

#### Physical Motivation

The Lateral Average Model (LAM) allows for modeling varying flow distributions from the vasculature to surrounding tissue. Exchanges occur both along each vessel segment in the vasculature graph and at the terminal nodes of the vessel. The amount of exchange is controlled by a drainage coefficient that may take a different value for each vessel segment. Along the vessel segment, the flow distribution depends on the difference between the pressure in the vessel segment and the tissue pressure that has been averaged laterally along the surface of the generalized cylinder that represents the section of the physical blood vessel with centerline equal to the vessel segment.

#### 3D Tissue Model Problem

Recall that *A*_*i*_, 1 ≤ *i* ≤ *N*_*out*_ are the terminal nodes of the network. In the 3D tissue domain, we solve for the pressure *p*_*t*_ that satisfies

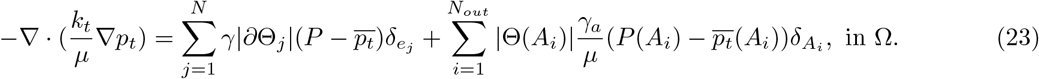

The Dirac measure 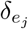 is concentrated on the line *e*_*j*_ and the Dirac measure 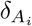 is concentrated at the point *A*_*i*_. The lateral average of *p*_*t*_ is denoted by 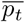 and defined by:

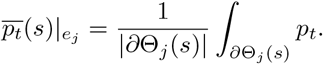

The boundary of the tissue Ω is divided into two parts ∂Ω_*N*_ and ∂Ω_*out*_. The terms in the right-hand side of (23) represent the coupling along the length of the vessel and at the terminal nodes respectively.

Homogeneous Neumann boundary conditions are imposed on ∂Ω_*N*_ and Robin boundary conditions are imposed on ∂Ω_*out*_. Let **n** denote the unit outward normal to Ω:

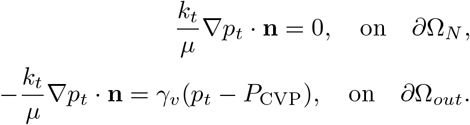

#### 1D Vasculature Model Problem

On each network edge *e*_*j*_, we solve for:

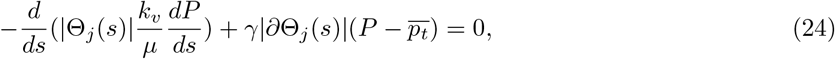

such that along each centerline Λ_*i*_, set

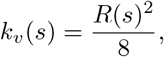

where *R*(*s*) is the local radius and with suitable conditions at the bifurcation nodes of the network, namely continuity of pressure and Kirchhoff conditions. The system is complemented with Dirichlet boundary condition at the inlet nodes, denoted by *B*_*i*_, and Neumann boundary conditions at the terminal nodes *A*_*i*_ that represent the coupling between the vessels and tissue:

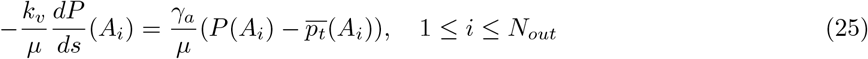

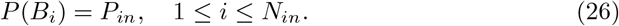

#### Discrete Model Problem

We partition the domain Ω into tetrahedra of size *h*_Ω_ and the network edges *e*_*j*_ into intervals of size *h*_*G*_. Let *V*_*h*_ be the continuous finite element space of order one over the partition of Ω and let *X*_*h*_ be the continuous finite element space of order one over the partition of *E*. Let *X*_*h*0_ be the subset of *X*_*h*_ of functions that vanish at the inlet nodes *B*_*i*_. The discrete solutions *p*_*h*_ ∈ *V*_*h*_ and *P*_*h*_ ∈ *X*_*h*_ satisfy Eq (26) and the following discrete variational problem: for any *v*_*h*_ ∈ *V*_*h*_ and *w*_*h*_ ∈ *X*_*h*0_,

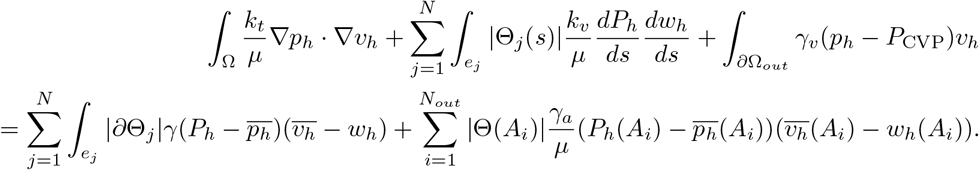

It is easy to show that there exists a unique solution (*p*_*h*_, *P*_*h*_) to the discrete problem.

### Using Clinical Data

#### Animal Protocol

This study was conducted at the University of Texas MD Anderson Cancer Center, Houston, TX, USA under an institutionally approved protocol (IACUC protocol number 00001478-RN03 Approved 8/20/2024 by MD Anderson Cancer Center Institutional Animal Care and Committee), using three outbred swines. The animals were purchased from Oakhill Genetics, Ewing Illinois. All methods were carried out in accordance with relevant guidelines and institution policy. All the methods are reported in accordance with ARRIVE guidelines for the reporting of animal experiments. The animals were acclimated and housed according to institutional policy. After induction and intubation, anesthesia was maintained with 2 % isoflurane, and supplemental oxygen was provided as needed. Buprenorphine was administered at 0.02 mg kg^−1^ intramuscularly for analgesia. The microcatheter was positioned proximally in a lobar artery. The injection of the reagent was done by hand in real time under direct fluoroscopic visualization and at a rate that allowed blood flow to carry the droplets forward and not to cause reflux. Thus, forward flow was initially at the same velocity as blood flow but gradually decreased until it stopped completely. The procedure was terminated when either the target volume (600 µL) was delivered or further infusion would cause reflux. Following each experimental procedure, animals recovered and were monitored until return to baseline activity and food intake. Euthanasia after completion of the study was performed via overdose of phenytoin and pentobarbital given intravenously while animals were under general anesthesia. An iodinated contrast medium (Visipaque 320; GE Healthcare, Milwaukee, WI) was used directly as a supplied as a contrast agent for CT scans.

#### Image Acquisition

Images of the animals were acquired using a 128-slice computed tomography (CT) system (SOMATOM Definition Edge; Siemens Healthineers, Forchheim, Germany). The CT scanner was part of a hybrid suite used in combination with an Artis Q angiography unit (Siemens Healthineers). Pretreatment/postcontrast and posttreatment/precontrast CT images were obtained. CT hepatic arteriography was performed by inserting a microcatheter into the common hepatic artery, injecting the contrast medium, and scanning after a suitable delay. All scans were acquired with a tube voltage of 120 kVp, rotation time of 0.5 s, pitch of 0.6, and 350 effective mA s. The resulting volumetric CT dose index for each scan was 23.45 mGy. Reconstructions were performed at a slice thickness and interval of 0.5 mm, display field of view of 420 mm, and corresponding in-plane pixel size of 0.8 mm.

#### Image Processing

Vessel segmentation was performed with a Hessian-based vesselness filter [**?**]. The parameter value thresholds for the blob structure, plate structure, and second-order structures were 0.5, 0.5, and 5.0, respectively, with a Gaussian blur radius of 2 mm. The vesselness filter was set at a threshold of 1 to generate the corresponding segmentation images. A 3D thinning algorithm was applied to extract the vessel centerline [**?**]. A sign distance transform was applied to computation of the radial distance from the segmentation boundary and the vessel centerline. Landmarks were manually placed on vessel bifurcations of pretreatment/postcontrast imaging and posttreatment/precontrast images. Depending on the visible anatomy in the images, 5-10 landmarks were placed on each image. Landmark-based registration was applied to alignment of the pretreatment and posttreatment images. Fig. 2 shows the centerline extraction of the hepatic artery from the arterial phase of CTHA.

**Fig 2.**
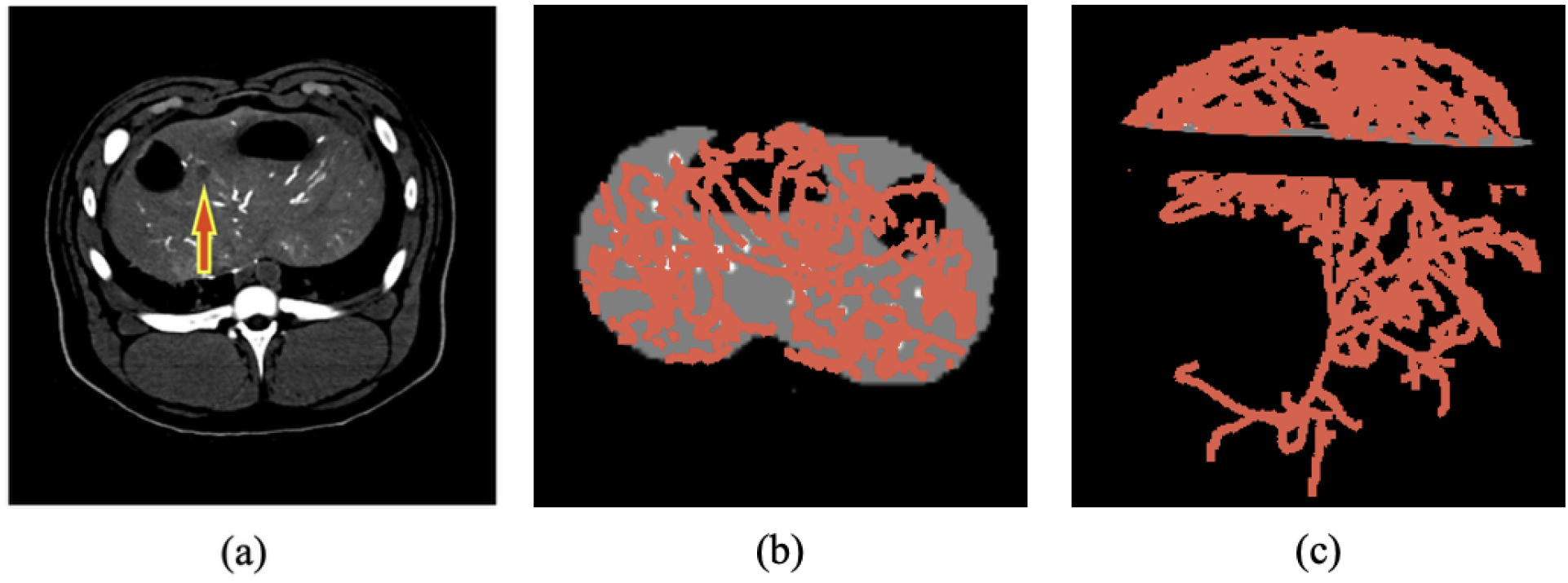
Protocol differences and timing of the CT acquisition change the enhancement of arterial vasculature and can impact image segmentation. (a) The arterial phase of a CTHA in a pig model is shown. The red arrow indicates the location of the target lesion. Our imaging protocols provide good contrast between the blood vessels and background liver. (b) Blood vessels are segmented using a Hessian-based vesselness filter. (c) A 3D model of the vasculature 1D centerlines is displayed with respect to the image.

The selected parameters for the Hessian-based vesselness filter balance anatomical specificity and imaging physics. The blob (*α* = 0.5) and plate (*β* = 0.5) thresholds prioritize tubular structures by penalizing eigenvalue ratios indicative of spherical or planar geometries, suppressing non-vascular features like tumors or imaging artifacts. These values align with Frangi’s recommendation (*α*/*β* ∈ [0.5, 2.0]) for vascular applications [**?**]. The second-order structure threshold (c = 5.0) amplifies responses for high-contrast vessels in CTHA, where iodinated agents create intense arterial signal against parenchyma. This compensates for the Frangi filter’s inherent sensitivity decay at larger scales. The 2 mm Gaussian blur (*σ*) corresponds to half the hepatic artery’s typical diameter (3–5 mm), optimizing Hessian responses while smoothing noise without oversmoothing bifurcations [**?**]. Combined, these parameters enhance medium-to-large hepatic arteries while rejecting smaller noise-induced pseudo-vessels and non-tubular anatomies, as validated in vascular segmentation benchmarks [**?**].

#### Converting an NifTi File to a 1D Vascular Network

The centerline extracted from the segmentation of the arterial phase of CTHA is stored in NifTi file format. The NIfTI file contains centerline voxels, information about the vessel radius, and ethiodized oil (Lipiodol) distribution in the vasculature. A Python script processes the NIfTI file to identify these centerline voxels. The script then employs a connectivity algorithm to construct the vascular tree as follows: 1. The Python script examines each voxel’s immediate neighborhood, considering all adjacent voxels (including diagonals) within a one-voxel distance. 2. Connected voxels are linked, and the length of each connection is calculated. 3. This process continues iteratively until all possible voxel connections are established. Because of limitations in imaging resolution or data quality, the result of this script may not be a single continuous structure. Instead, multiple disconnected trees may form with gaps in the data that prevent direct connections. In such cases, the largest continuous tree (i.e., the one with the most connected voxels) is selected for further analysis provided it accurately represents the hepatic artery. This approach ensures that the simulations are based on the most complete and representative vascular structure available from each imaging datasets while maintaining the integrity of the analysis by excluding inadequate reconstructions.

Fig. 3.(a) shows the entire segmented hepatic artery and liver parenchyma. Due to the computational limit, a decision was made to focus only on the treatment region i.e. from the point of injection till the last observed lipiodol post treatment. The entire liver data is of size 435.2 mm × 435.2 mm × 161 mm. The computational control volume (denoted by CV) is shown in Fig. 3.(b); the selected CV size is 103.7 mm × 53.5 mm × 52.5 mm. The CT scans used for this data did not have the hepatic veins visible and we could not segment them for this simulation. An assumption was made that the probability of a draining hepatic vein to exist is higher in the upper region of the control volume. Based on that assumption a sink region was identified which was used to impose the Robin boundary condition as shown in Fig. 3.(c).

**Fig 3.**
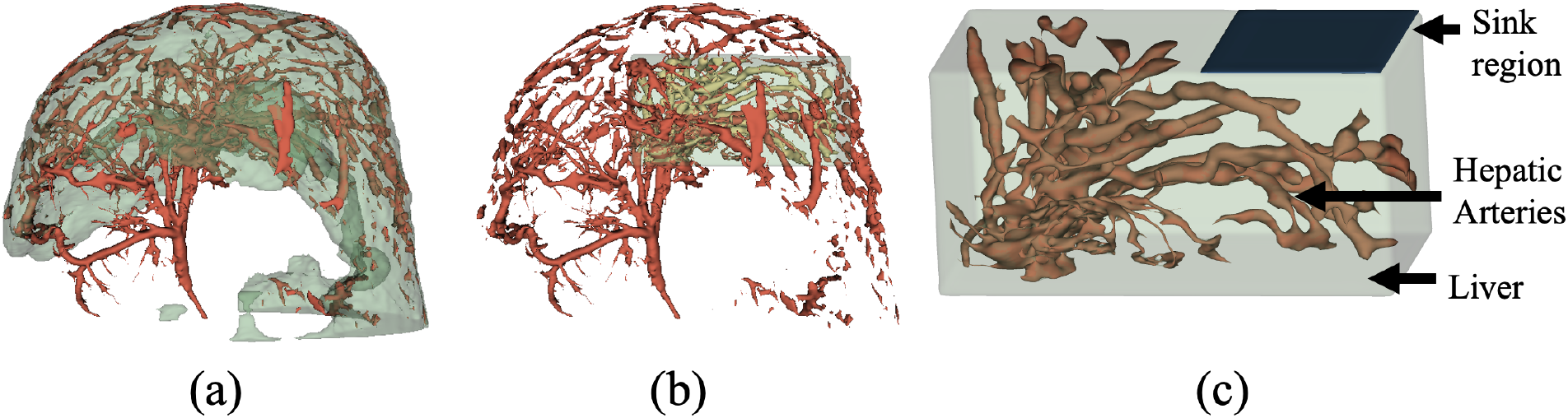
(a) Entire segmented hepatic artery and liver parenchyma of swine obtained from a CT scan. (b) The simulation domain focused only on the region treated during the animal study (c) Focused view of the clinical demo domain used for this study with liver, hepatic artery and sink region labeled.

The CV was used to compare global behavior of flow between the two models. To further analyze any local variations in the CV, two distinct regions of influence (ROIs) were determined. These ROIs are shown in Fig. 4, and the specific ROIs will be referred to as ROI 1 and ROI 2, respectively.

**Fig 4.**
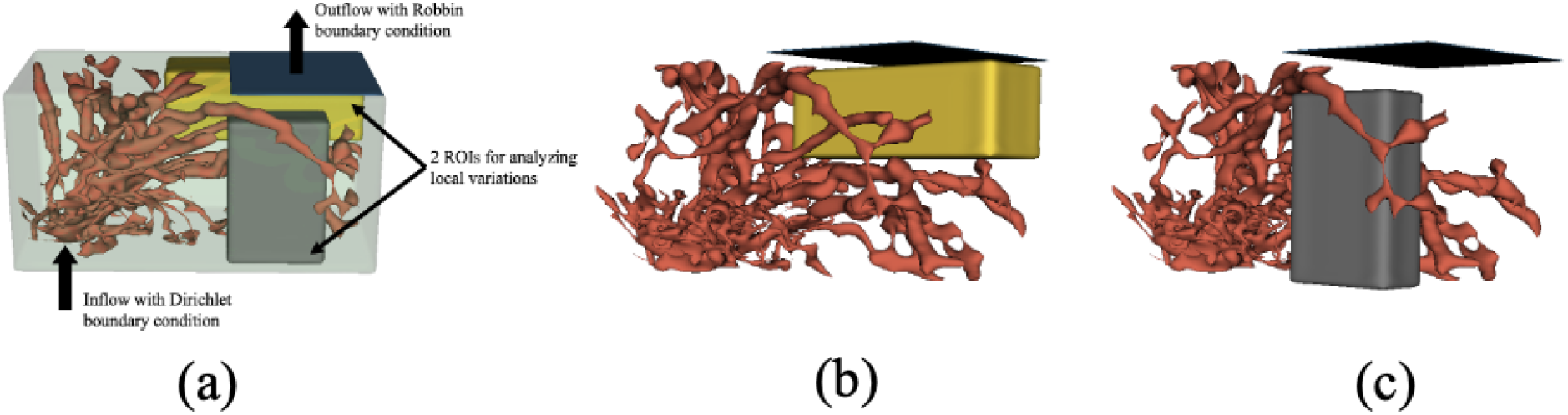
(a) The ROIs identified for analyzing local variations in two models. (b) ROI 1 (c) ROI 2

For this animal study analysis, the parameters used for simulation are given in Table 1. For a comprehensive analysis the values of inlet pressure (*P*_in_) were varied in the range of 80 mm Hg to 100 mm Hg, which is the range of swine blood pressure on arterial side.

**Table 1.**
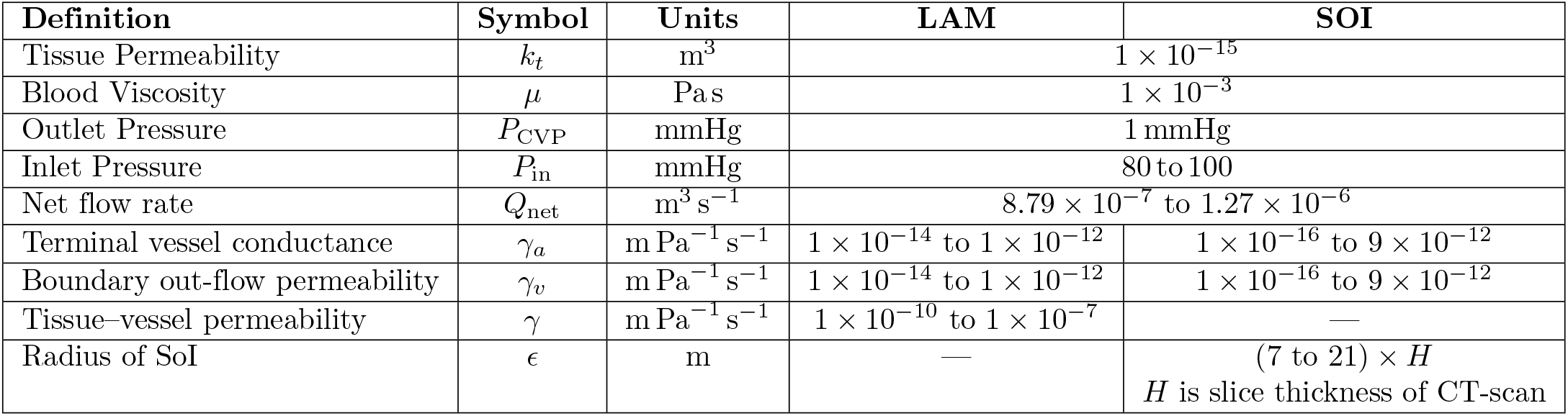
Swine liver simulation parameters.

## Results

### Comparison between the two models on benchmark model

The set-up of the problem is given in Fig. 5.

**Fig 5.**
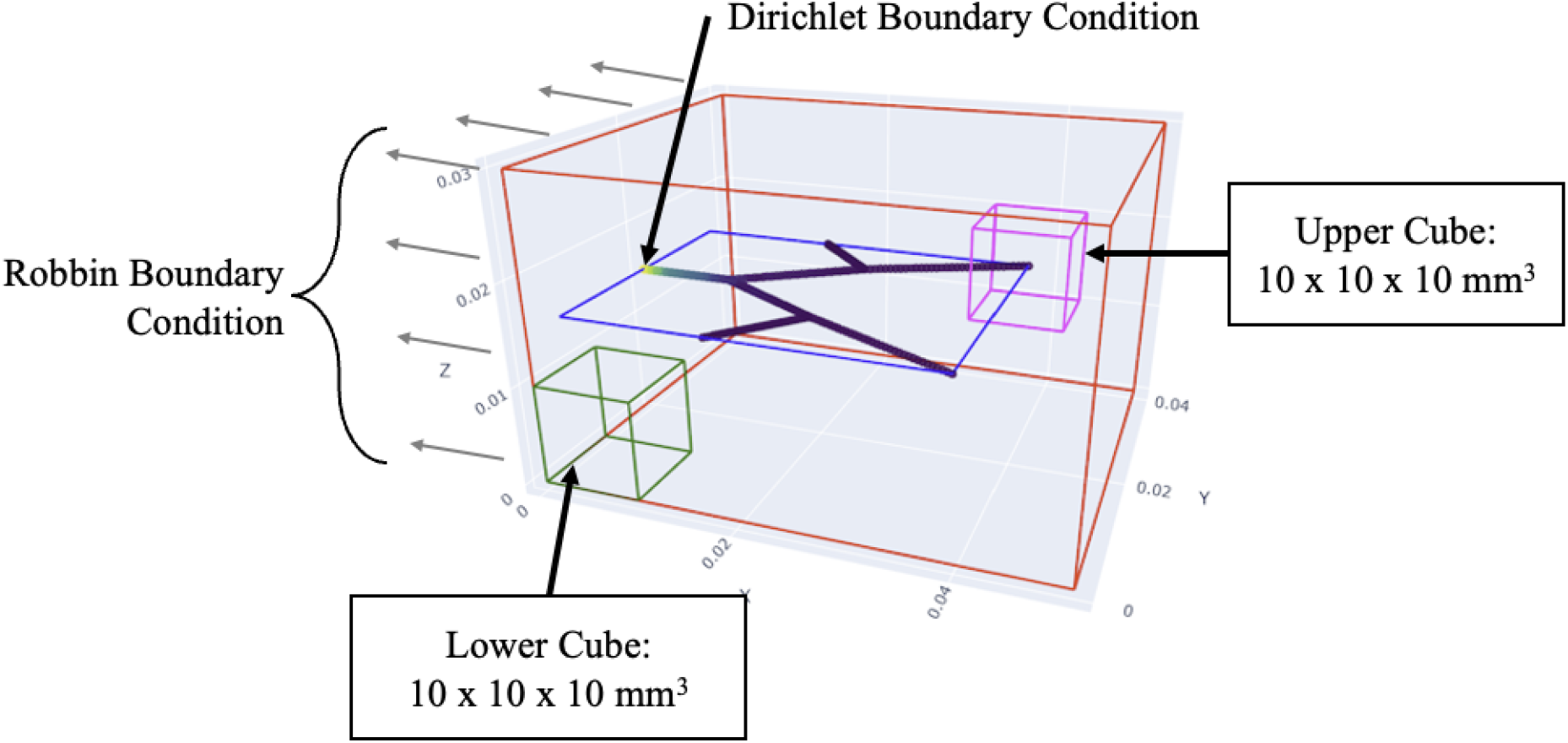
Domain and parameters.

**Fig 6.**
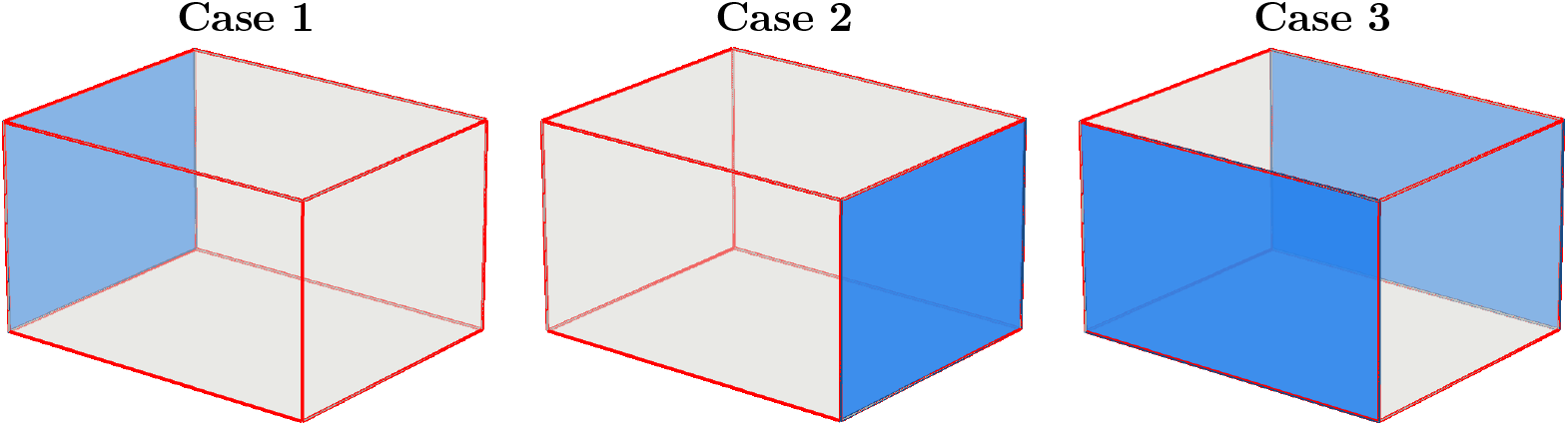
Three different sink boundary configurations: a sink at the *x* = 0 mm face of the 3D domain (Case 1), a sink at the *x* = 50 mm face (Case 2), and sinks at the *y* = 0 mm and *y* = 40 mm faces (Case 3).

The computational domain is a box of dimensions 50 × 40 × 30 mm^3^ representing the 3D tissue. A simple network of vessels is placed in the horizontal plane at height 15 mm. Coordinates of each graph node and radii of each vessel are given in S1 Fig. We will evaluate the LAM and SOI model by focusing on two spatially distinct cubic regions of interest (ROIs), each measuring 10 × 10 × 10 mm^3^, representing potential tumor locations within a tissue domain perfused by a 1D vascular network. The upper cube is positioned in the interior of the domain where multiple vessel terminals are present, making it a primary recipient of 1D-to-3D coupling in the SOI model. The lower cube is located near the case 1 sink (outlet) boundary condition, geometrically distant from most vessel terminals. The 1D vessel network traverses through the domain with vessels terminating primarily in the upper and middle regions. The domain is subjected to mixed boundary conditions: Dirichlet boundary conditions prescribe pressure at the boundaries, while Robin boundary conditions govern the lateral coupling between the 1D vessels and 3D tissue in the LAM. This experimental configuration enables systematic investigation of how coupling mechanism choice (LAM versus SOI) and parameter selection (*γ* versus *ε*) influence regional blood flow distribution based on spatial proximity to coupling sites, a critical consideration for predicting drug delivery, hypoxia patterns, and treatment response in heterogeneous tumor microenvironments.

#### Case 1

Fig. 7 compares the midplane pressure field produced by LAM and SOI when the venous sink is collocated with the arterial inlet face of the network. In the LAM panel (a) the vessel network appears as a thin curve carrying the 1D pressure *P*, while the surrounding tissue pressure *p*_*t*_, which is solved from Eq. (23) with distributed lateral exchange along the entire centerline E – is elevated throughout the central region of the slice and decays only in a narrow band adjacent to the Robin outflow boundary ∂Ω_*out*_ at *x* = 0.

**Fig 7.**
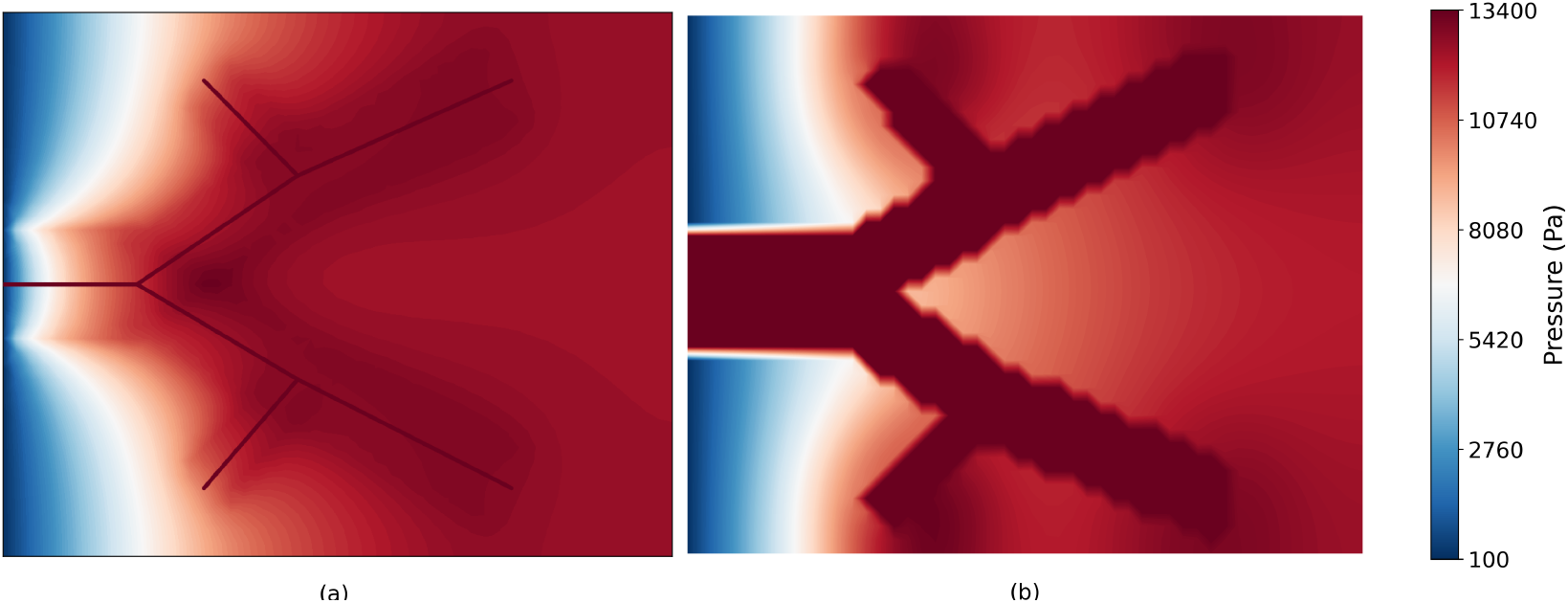
Midsliceplane pressure profiles for sink boundaries placed at the left face (*x* = 0) via (a) LAM with *γ*_*a*_ = 0, *γ*_*v*_ = 10^−7^, *γ* = 5 × 10^−10^, and (b) SOI with *γ*_*a*_ = *γ*_*v*_ = 2 × 10^−10^ and *ε* = 12mm.

In the SOI panel (b) the same network appears as a thick connected red region: with *ε* = 12 mm, the spheres *B*_*ε*_(*A*_*k*_) around the (closely spaced) arterial terminals overlap heavily. Substantial pressure drop happens between the terminal of the segmented blood vessel and the tissue (*P* ≫ *p*_*t*_) and is controlled by *γ*_*a*_ parameter in SOI. Outside the sphere union, *p*_*t*_ decays diffusively toward *P*_CVP_ within the sink region Ω_sink_ near *x* = 0. Because the inlet face and the drainage region coincide, the global pressure drop across the slice is small in both models: most of the leaked fluid finds the sink before traversing the bulk of the tissue.

Fig. 8 shows the global net flow rate (*Q*_*net*_) through the 3D tissue domain (panel a) together with the net outflow leaving the lower (panel b) and upper (panel c) ROIs, plotted against the LAM permeability *γ* (blue) and the SOI sphere radius *ε* (orange).

**Fig 8.**
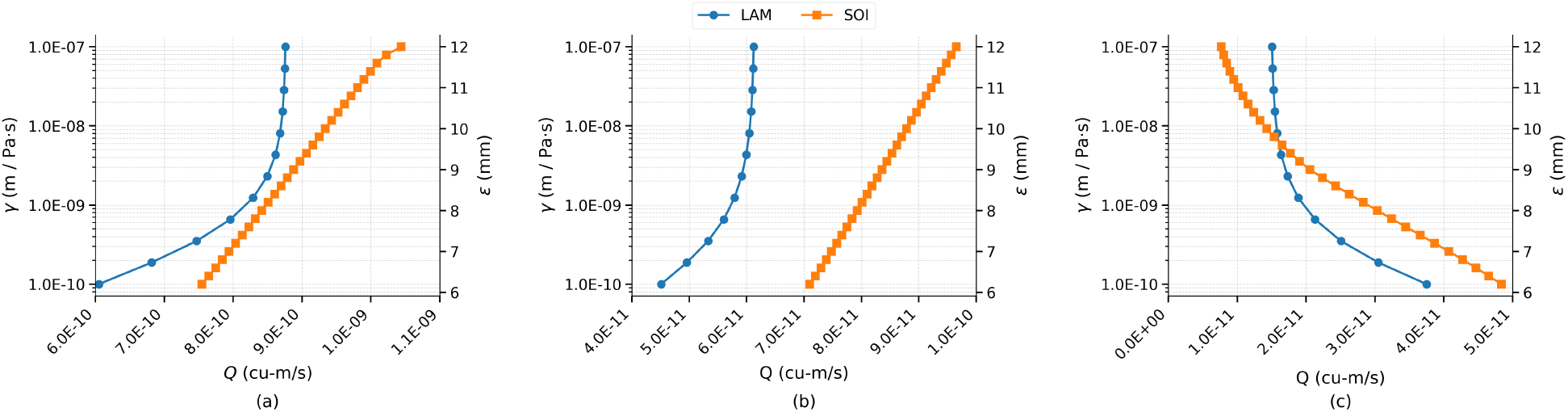
Case 1 comparison of two models. (a) Variation of total volumetric flow rate in the entire CV as *γ* and *ε* are varied, respectively. (b) Variation of the outflow volumetric rate from lower ROI in two models. (c) Variation of the outflow volumetric rate from upper ROI in two models.

In panel (a), the two coupling formulations exhibit distinct parametric responses. The LAM curve rises as *γ* increases from 10^−10^ to ~10^−9^ m/Pa·s and then asymptotes toward a plateau near 8.7 × 10^−10^ m^3^/s for *γ* ≳ 10^−8^. This saturation indicates that once the lateral exchange is sufficiently permeable, the rate-limiting step shifts from the 1D–3D coupling itself to bulk transport in the 3D compartment. In contrast, the SOI net flow rate grows almost linearly with *ε* over the tested range *ε* ∈ [5.8, 12] mm, without saturation. Despite this qualitative difference, both models converge to comparable magnitudes of *Q*_*net*_ at their respective upper parameter limits, indicating that equivalent global perfusion can be achieved through either coupling strategy with appropriate parameter choice.

In panel (b), the lower ROI shows a strictly positive net outflow for both models, confirming that this region–geometrically close to the *x* = 0 outlet– acts as a local sink that drains interstitial fluid toward the boundary. The LAM outflow increases from ~4.5 × 10^−11^ at *γ* = 10^−10^ to a plateau near 6.5 × 10^−11^ m^3^/s for *γ* ≳ 10^−8^, mirroring the saturation observed in *Q*_*net*_. The SOI outflow increases monotonically and roughly linearly from ~7 × 10^−11^ at *ε* = 5.8 mm to ~1 × 10^−10^ m^3^/s at *ε* = 12 mm. The two models produce a flow of same order of magnitude in this region.

Panel (c), the upper ROI, exhibits a markedly different and counterintuitive response. This region contains a section of the terminal vessel branches, so the vessel network deposits mass directly within it; in steady state, that mass must leave the ROI through its bounding faces, giving a strictly positive net outflow. However, the outflow *decreases* as either coupling parameter is increased: LAM outflow drops from ~ 3.7 × 10^−11^ at *γ* = 10^−10^ to ~ 1.5 × 10^−11^ m^3^/s at *γ* = 10^−7^, and SOI outflow drops from ~ 4.8 × 10^−11^ at *ε* = 5.8 mm to ~ 1.1 × 10^−11^ m^3^/s at *ε* = 12 mm. Two effects contribute to this trend, that is *opposite* to that of *Q*_*net*_. First, as the coupling becomes spatially more diffuse–vessel mass exchanged along longer segments (LAM) or distributed over larger spheres (SOI)–a growing fraction of the source is deposited *outside* the upper ROI envelope, so less of the total source must transit the ROI’s bounding faces. Second, because the sink boundary lies on the same face from which the vessel network feeds, there is an effective short-circuit between the vessel inlet and the *x* = 0 outlet: stronger coupling causes more flow to leak from the vessel early in its path and drain directly to the nearby sink, bypassing the deeper terminal region that lies within the upper ROI.

#### Case 2

Fig. 9 shows the qualitatively different regime that emerges when the Robin/sink boundary is moved to the face opposite the vessel inlet. Both panels now display a clean, almost one-dimensional gradient from red on the left (high pressure, where the network feeds in and exchanges with tissue) to deep blue on the right (low pressure at the drainage face). In LAM the gradient is smooth across the entire tissue because the distributed line-source exchange along ℰ keeps *p*_*t*_ elevated continuously over the network’s footprint; the result is a wide red plateau that fades through orange before reaching the Robin boundary.

**Fig 9.**
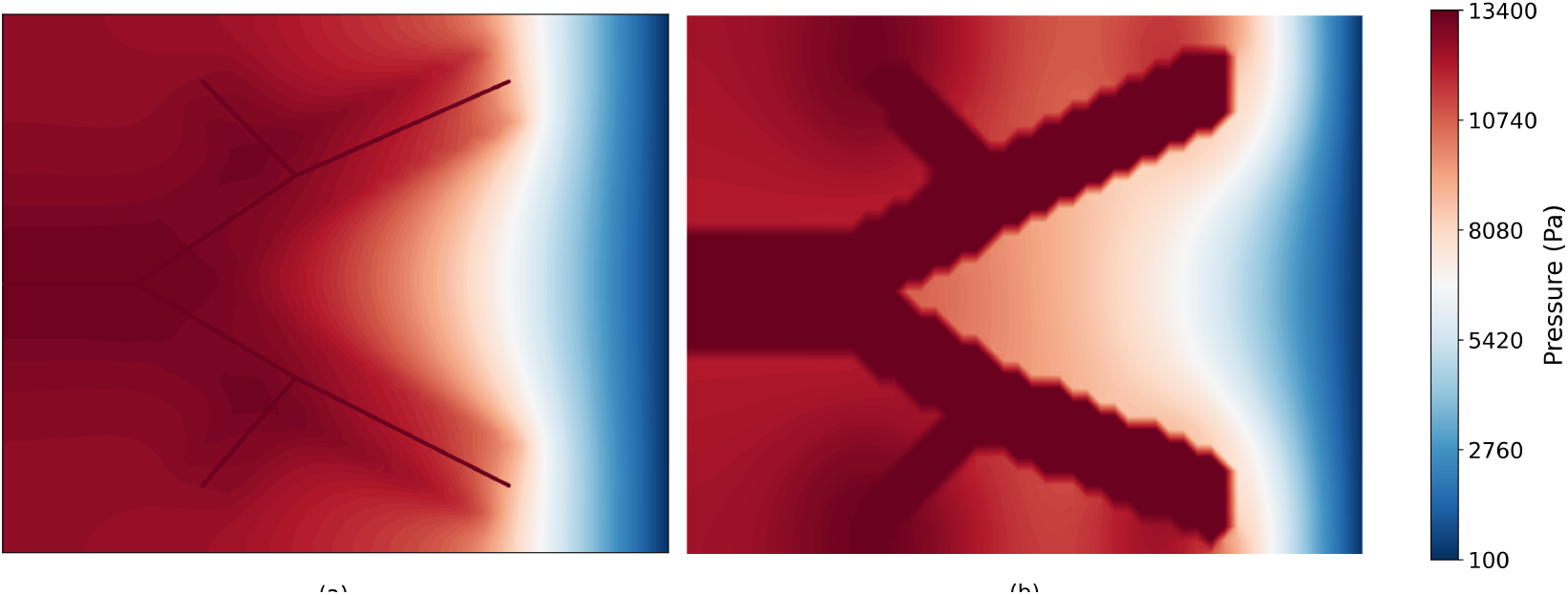
Midsliceplane pressure profiles for sink boundaries placed at the left face (*x* = 0.05) via (a) LAM with *γ*_*a*_ = 0, *γ*_*v*_ = 10^−7^, *γ* = 5 × 10^−10^, and (b) SOI with *γ*_*a*_ = *γ*_*v*_ = 2 × 10^−10^ and *ε* = 12mm.

In SOI the high-pressure region is more sharply delimited to the union of terminal spheres – outside that union the only mechanism elevating *p*_*t*_ is diffusion from the spheres, so the orange-to-blue transition occupies a larger fraction of the slice. In both cases the inlet face and the sink face now sit on opposite sides of the network, so the lateral exchange (LAM) and the terminal sphere sources (SOI) feed flow that *must* traverse the tissue toward *x* = 50 mm. The upper ROI, which lies on this through-route, is therefore on the natural drainage path, and any extra flow released by raising *γ* or *ε* is funnelled through it – explaining the increasing upper-ROI outflow in LAM and the rise-then-fall behaviour in SOI seen in Fig. 10(c). The visibly larger pressure drop across the slice compared to Case 1 is also the geometric reason for the much higher *Q*_*net*_. We also note that for the SOI method, the blood vessels retain a 3D volume.

**Fig 10.**
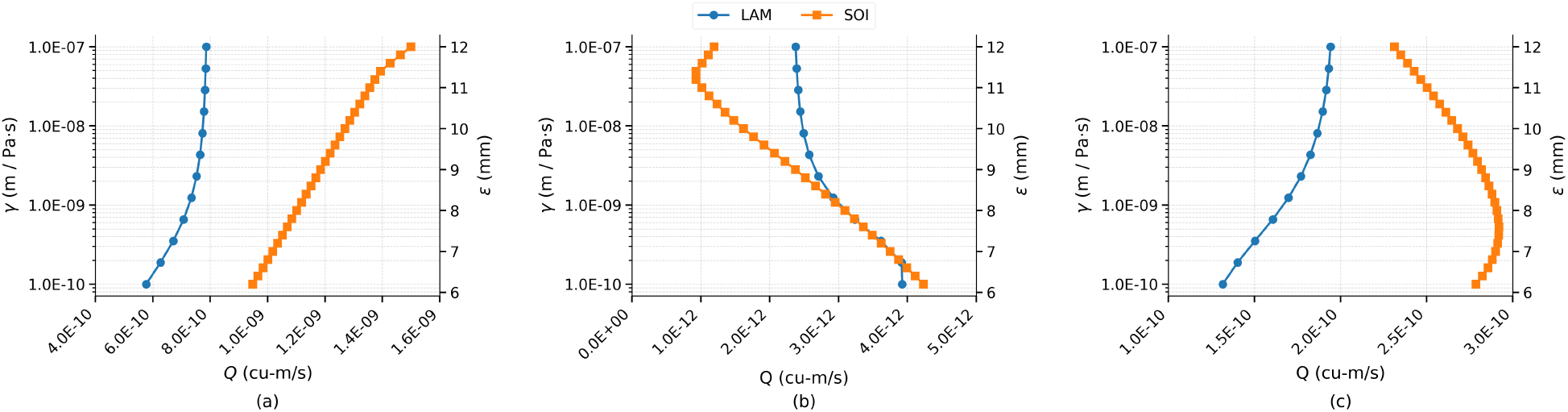
Case 2 comparison of two models. (a) Variation of total volumetric flow rate in the entire domain as *γ* and *ε* are varied, respectively. (b) Variation of the outflow volumetric rate from lower ROI in two models. (c) Variation of the outflow volumetric rate from upper ROI in two models.

Relocating the sink boundary to the *x* = 50 mm face fundamentally changes the perfusion regime relative to Case 1 by aligning the global pressure gradient with the natural direction of vessel-to-tissue flow. In Fig. 10(a), the LAM *Q*_*net*_ rises with *γ* to a plateau near 8.2 × 10^−10^ m^3^/s, comparable in magnitude to the Case 1 plateau, while the SOI *Q*_*net*_ grows from ~ 7.5 × 10^−10^ m^3^/s at *ε* = 5.8 mm to ~ 1.5 × 10^−9^ m^3^/s at *ε* = 12 mm without saturation. In panel (b), the lower ROI shows outflows roughly an order of magnitude smaller than Case 1 (~1–4 × 10^−12^ m^3^/s), with a weak, slightly decreasing dependence on the coupling parameter; because the sink boundaries lie transverse to the dominant vessel orientation, the lower cube is no longer on the natural drainage path and instead acts essentially as a flow-through region with small net imbalance across its faces. The upper ROI (panel c) shows the LAM outflow rising monotonically from ~1.3 × 10^−10^ m^3^/s at *γ* = 10^−10^ to a plateau near 1.9 × 10^−10^ m^3^/s for *γ* ≳ 10^−8^, while the SOI outflow rises from ~2.5 × 10^−10^ m^3^/s at *ε* = 5.8 mm to a peak of approximately 2.9 × 10^−10^ m^3^/s near *ε* ≈ 7.5 mm, then decreases to ~2.5 × 10^−10^ m^3^/s at *ε* = 12 mm. The peak corresponds to the geometric crossover at which the sphere of influence around each terminal first exceeds the distance from the terminal to the nearest ROI face (roughly the half-width ~5 mm): for smaller *ε* the sphere fits entirely within the ROI so the entire source must leave through its faces, whereas for larger *ε* an increasing fraction of the sphere protrudes outside the ROI and that mass is delivered directly to the surrounding tissue.

#### Case 3

Fig. 11 shows the pressure contour map for Case 3 for sink boundaries placed at the two side faces (*y* ∈ [0, 0.04]). For LAM the tissue-vessel permeability was *γ* = 5 × 10^−10^ with *γ*_*a*_ = 0, *γ*_*v*_ = 10^−7^ and for SOI the radius *ε* = 12mm with *γ*_*a*_ = *γ*_*v*_ = 2 × 10^−10^. With sinks now imposed simultaneously on the *y* = 0 and *y* = 40 mm faces, the domain has two parallel transverse drainage pathways. The Robin boundary condition in LAM now applies on two faces, ∂Ω_*out*_ = {*y* = 0} ∪ {*y* = 40}, and in SOI the sink region Ω_sink_ occupies slabs along both transverse faces. Robin boundary condition in LAM now applies on two faces, ∂Ω_*out*_ = {*y* = 0} ∪ {*y* = 40}, and in SOI the sink region Ω_sink_ occupies slabs along both transverse faces. Compared with Case 2, the high-pressure core in both panels is *more confined* and the pressure drop is *more localised* near the sink-facing boundaries: the bulk of the slice containing the network remains close to vessel pressure (large red interior in both LAM and SOI), and the steep decay is concentrated in narrow boundary layers adjacent to the two transverse drainage faces. Mathematically this reflects that the distance from any source point to the nearest sink is shorter (and the available drainage cross-section larger) than in the single-face configurations, so diffusion needs less travel to remove the deposited mass. The lower ROI lies near the *y*-centerline of the domain, away from both drainage faces, so it sits in the high-pressure plateau and is no longer on a strong gradient – consistent with its order-of-magnitude smaller net outflow in Fig. 12(b). The upper ROI, which contains the terminal nodes themselves, sits inside the red core of both contour maps and registers the largest upper-ROI outflow of the three cases under SOI, with the same non-monotonic geometric peak at *ε* ≈ 7.5 mm that arises when the sphere of influence first exceeds the terminal-to-ROI-face distance.

**Fig 11.**
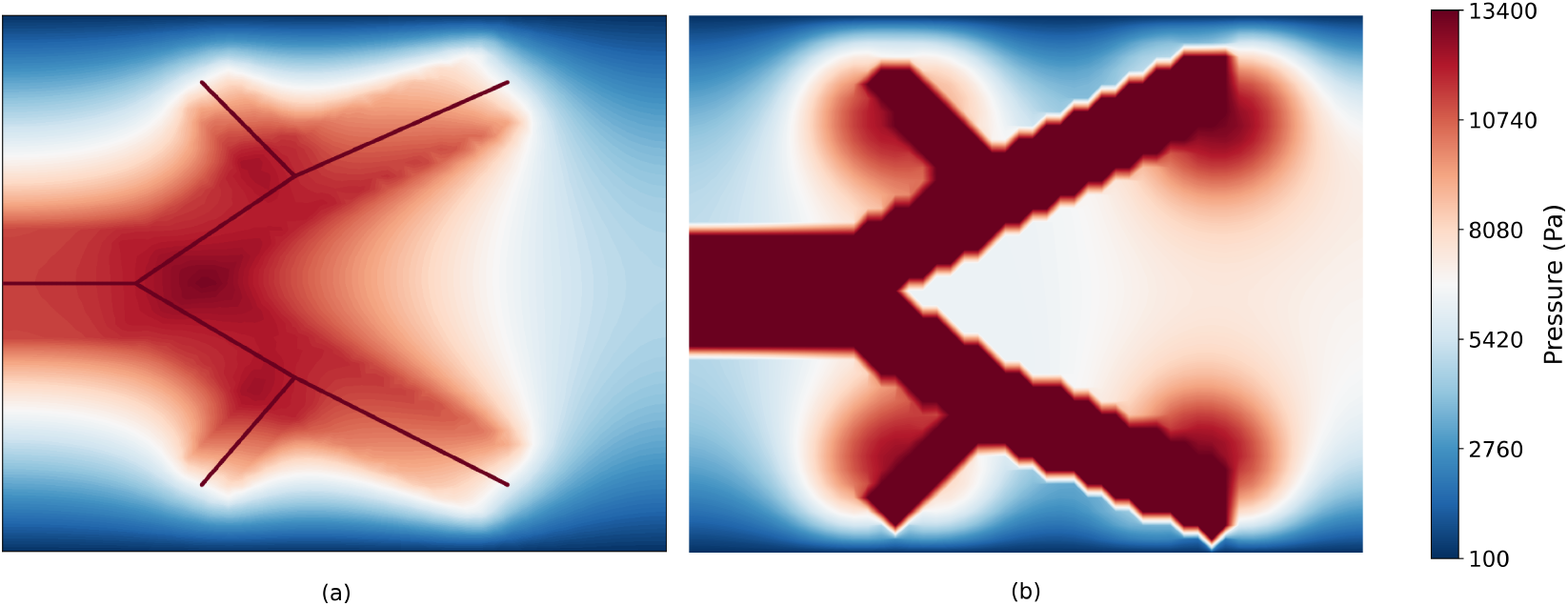
Midsliceplane pressure profiles for sink boundaries placed at the the two side faces (*y* ∈ [0, 0.04]) via (a) LAM with *γ*_*a*_ = 0, *γ*_*v*_ = 10^−7^, *γ* = 5 × 10^−10^, and (b) SOI with *γ*_*a*_ = *γ*_*v*_ = 2 × 10^−10^ and *ε* = 12mm.

**Fig 12.**
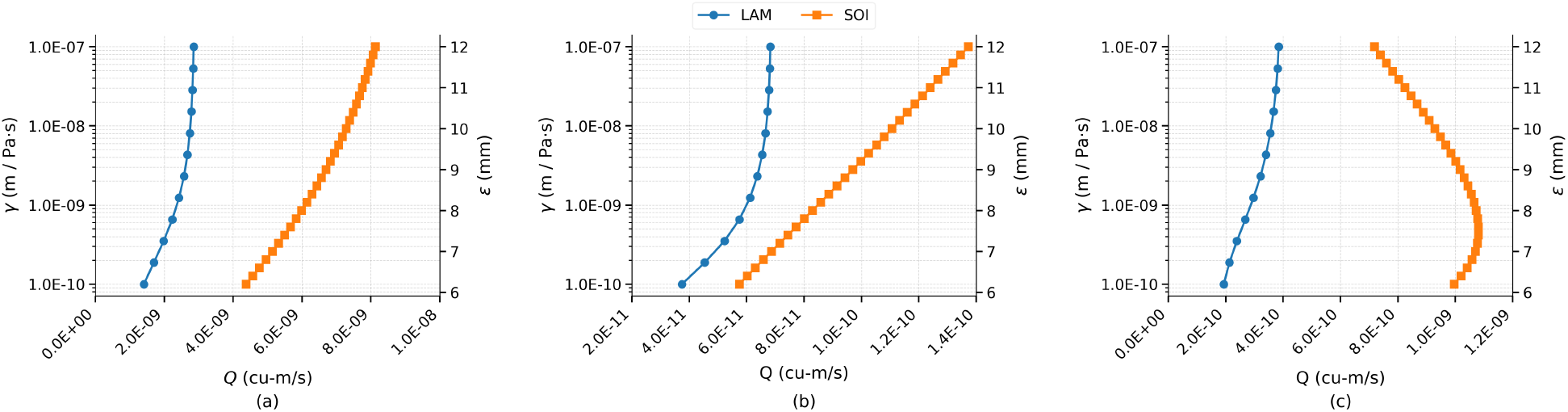
Case 3 comparison of two models. (a) Variation of total volumetric flow rate (*Q*_*net*_) in the entire domain as *γ* and *ε* are varied, respectively. (b) Variation of the outflow volumetric rate from lower ROI in two models. (c) Variation of the outflow volumetric rate from upper ROI in two models.

In Fig. 12(a), the LAM *Q*_*net*_ rises with *γ* to a plateau near 2.8 × 10^−9^ m^3^/s — roughly three times the Case 1 plateau — while the SOI *Q*_*net*_ increases across the entire *ε* range, from ~4 × 10^−9^ m^3^/s at *ε* = 5.8 mm to ~8 × 10^−9^ m^3^/s at *ε* = 12 mm, without saturation. The two-to-three-fold gap between SOI and LAM magnitudes reflects the SOI’s greater sensitivity to source distribution when the sink lies far from the vessel inlets. In panel (b), the lower ROI shows monotonically increasing net outflow for both models, with LAM outflow saturating near 6.7 × 10^−11^ m^3^/s for *γ* ≳ 10^−8^ and SOI outflow rising linearly from ~ 6 × 10^−11^ to ~ 1 × 10^−10^ m^3^/s. The upper ROI (panel c) shows the LAM outflow rising with *γ* from ~ 2 × 10^−10^ to a plateau near 4 × 10^−10^ m^3^/s — an order of magnitude larger than in Case 1 — while the SOI response is non-monotonic: outflow rises from ~1 × 10^−9^ m^3^/s at *ε* = 5.8 mm to a peak of approximately 1.1 × 10^−9^ m^3^/s near *ε* ≈ 7.5 mm, then decreases back toward ~7 × 10^−10^ m^3^/s at *ε* = 12 mm. The location of the SOI peak at the same *ε* ≈ 7.5 mm as in Case 2 supports a geometric interpretation: the optimal sphere radius for maximising upper-ROI outflow is set by the distance from the terminal locations to the ROI faces, independent of the sink configuration. The overall picture is that LAM and SOI agree on the qualitative direction of upper-ROI flow response whenever the sink is well separated from the vessel inlets (Cases 2 and 3), but quantitatively the SOI delivers two-to-three times more flow through the upper ROI and does so non-monotonically, so the choice of coupling parameter has substantially greater consequences for local perfusion predictions than for global ones.

### Animal Study Analysis

The two coupling models are now evaluated on the physiologically realistic vascular geometry described in the *Using Clinical Data* section. The hepatic arterial network reconstructed from CT hepatic arteriography of the swine liver (Fig. 2.b and Fig. 2.c) provides a substantially more complex coupling topology than the benchmark network: the 1D vascular tree contains multiple branching generations resolved from the point of microcatheter injection to the distal terminals, embedded within a 103.7 × 53.5 × 52.5 mm^3^ tissue control volume (Fig. 3.c). The venous drainage geometry, which could not be resolved from the available CT imaging data, is represented by the Robin boundary condition imposed on the anatomically motivated sink region identified in the upper portion of the control volume (Fig. 3.c). Both LAM and SOI are applied to this control volume with the simulation parameters listed in Table 1, and inlet pressure *P*_in_ is varied over the physiological range of swine arterial pressure (80–100 mmHg) to assess model sensitivity under realistic hemodynamic boundary conditions. Global net flow rate *Q*_net_ and regional perfusion within the two identified ROIs (Fig. 4) are used as the primary comparison metrics, following the same framework established in the benchmark analysis.

Fig. 13 compares the global net flow rate *Q*_*net*_ produced by the two coupling formulations on the realistic vascular network, normalized onto a common, problem-independent scale

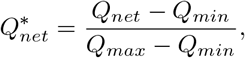

with *Q*_*min*_ = 8.79 × 10^−7^ m^3^/s and *Q*_*max*_ = 1.27 × 10^−6^ m^3^/s, and against a dimensionless sphere radius *ε*^∗∗^ = *ε/H* (*H* is the slice thickness of CT-scan) for SOI (left axis) paired with the LAM lateral permeability *γ* on a logarithmic scale (right axis). For each (*ε*^∗∗^, *γ*) row, the blue marker and shaded band give the SOI mean and the min–max envelope over the auxiliary couplings (*γ*_*a*_, *γ*_*v*_), while the red marker and band give the corresponding LAM mean and its min–max envelope over *γ*_*a*_, *γ*_*v*_. Table 1 shows the range of these parameters. Panels (a) and (b) repeat the analysis at inlet pressures *P*_*in*_ = 80 and 90 mmHg respectively. The vertical dashed lines at 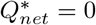 and 1 mark the reference window, and the grey regions identify predictions that fall outside it.

**Fig 13.**
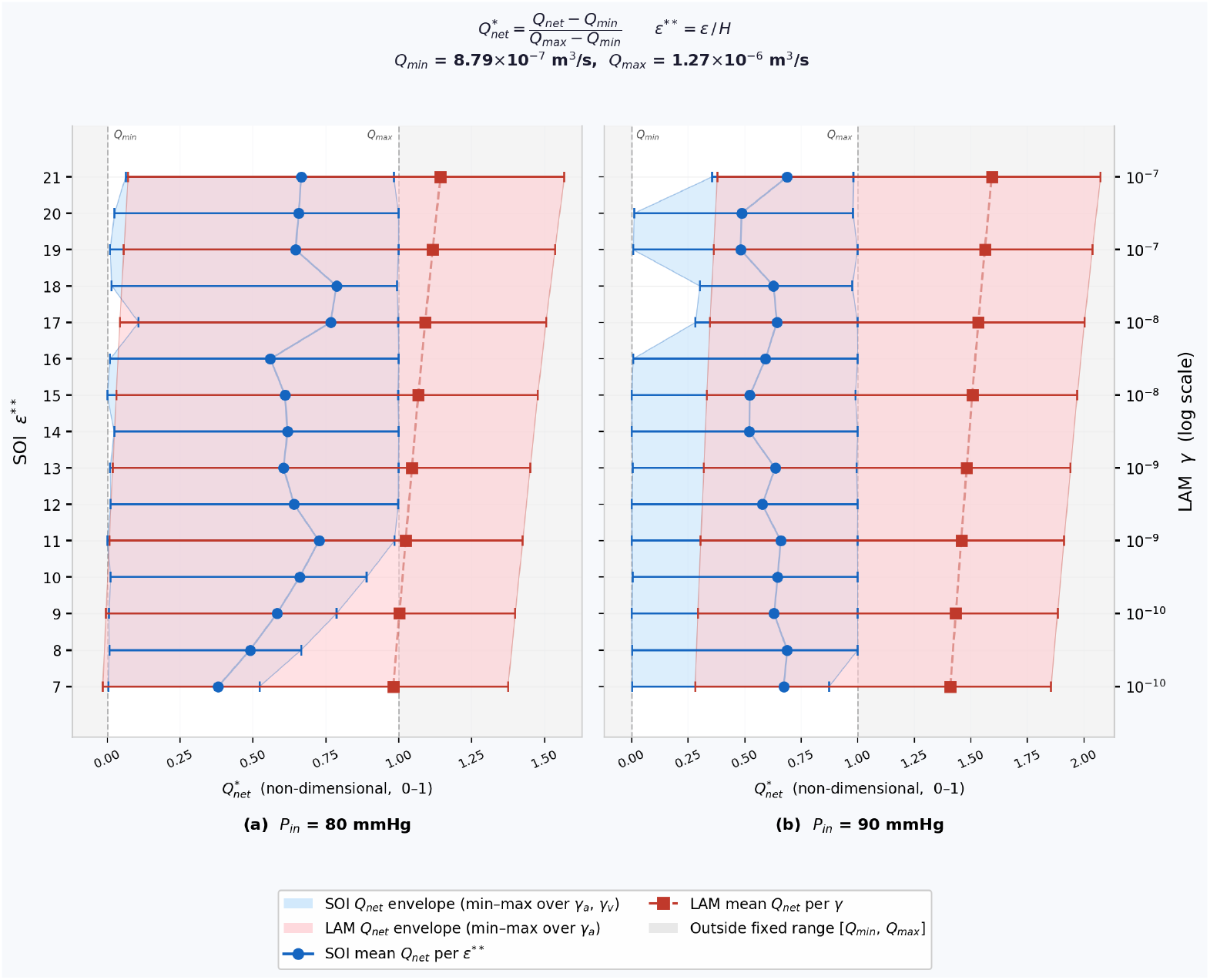
Normalized global net flow rate 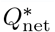 predicted by LAM and SOI on the swine liver control volume at inlet pressures (a) *P*_in_ = 80 mmHg and (b) *P*_in_ = 90 mmHg. The normalization 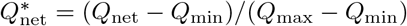 maps the physiological reference window [*Q*_min_, *Q*_max_] = [8.79 × 10^−7^, 1.27 × 10^−6^] m^3^s^−1^ to [0, 1] (vertical dashed lines); grey shading identifies predictions outside this window. For SOI (blue), each marker gives the mean 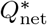 over the auxiliary coupling parameters (*γ*_*a*_, *γ*_*v*_) at a fixed dimensionless sphere radius *ε*^∗∗^ = *ε/H* (left axis, *H* = CT slice thickness), and the shaded band spans the corresponding min–max envelope. For LAM (red), each marker gives the mean 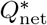 over *γ*_*a*_ at a fixed tissue–vessel permeability *γ* (right axis, logarithmic scale), with the shaded band spanning the min–max envelope over *γ*_*a*_. SOI predictions cluster within the physiological reference window across the full *ε*^∗∗^ range, while LAM predictions consistently exceed *Q*_max_ and exhibit a wider parametric spread driven primarily by sensitivity to *γ*_*a*_ rather than *γ*.

A first observation is that the two formulations sit on *systematically different* parts of the normalised axis. The LAM mean clusters tightly around 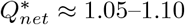 at *P*_*in*_ = 80 mmHg and ≈ 1.40–1.55 at *P*_*in*_ = 90 mmHg, placing it at or beyond the upper reference *Q*_*max*_ for essentially every value of *γ* tested. The near-invariance of the red mean across the four decades of *γ* is a direct manifestation of the saturation behaviour identified on the benchmark problem in Figs. 8–12(a): once *γ* is large enough that the lateral exchange ceases to be rate-limiting, the LAM net flow rate is set by the bulk tissue conductivity and the boundary configuration alone, so varying *γ* produces almost no movement along the 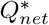 axis. The width of the red envelope, on the other hand, shows that LAM remains quite sensitive to *γ*_*a*_ while remaining relatively robust to variations in *γ*_*v*_ : the min–max band spans roughly 0.5–1.6 in panel a and 0.9–2.0 in panel b, indicating that the terminal-conductance parameter – not the distributed lateral permeability – is the dominant control on LAM perfusion at these parameter ranges.

The SOI predictions, by contrast, sit comfortably inside the reference window in both panels: the blue mean ranges over 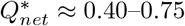 at *P*_*in*_ = 80 mmHg and 0.50–0.70 at 90 mmHg, with a mild and slightly non-monotonic dependence on *ε*^∗∗^ (a modest dip near *ε*^∗∗^ ≈ 12–16 followed by a recovery at the largest radii). The SOI envelope – min–max over both *γ*_*a*_ and *γ*_*v*_ – is markedly narrower than the LAM envelope, tightening further as *ε*^∗∗^ increases: at large radii the sphere of influence already covers enough tissue volume that the terminal conductance *γ*_*a*_ no longer strongly modulates the total source delivered, so the spread collapses toward the mean. This is the *opposite* of what we see in LAM, where larger *γ* does not narrow the *γ*_*a*_ envelope.

Comparing across the two inlet pressures, raising *P*_*in*_ from 80 to 90 mmHg shifts both formulations to higher 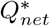 as expected from the linear pressure-driven character of the Stokes/Darcy system, but the shift is much larger for LAM than for SOI (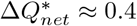 versus ≈ 0.05). Together these observations highlight a quantitative tension between the two models: under the same physiological constraints, LAM tends to over-predict perfusion relative to the prescribed reference window while SOI under-predicts it slightly but stays within physiological bounds. The choice between them therefore matters not only qualitatively – for the spatial distribution of flow, as documented in the benchmark-problem ROI analysis – but also quantitatively for total tissue perfusion, with *γ*_*a*_ in LAM and *ε*^∗∗^ in SOI emerging as the parameters that most directly control the predicted operating point.

## Discussion

This study presents a systematic quantitative comparison of the Lateral Average Model (LAM) and the Sphere of Influence (SOI) coupling paradigms for 3D-1D vascular-tissue flow simulations, revealing fundamental differences in how these formulations translate coupling parameters into hemodynamic predictions. Using a controlled computational framework with identical boundary conditions and vascular network geometry across both models, we isolated the effects of coupling mechanism from confounding numerical or geometric factors. Our results demonstrate that while both models can produce broadly comparable total domain flow rates under specific conditions, they exhibit markedly different parameter sensitivities, spatial redistribution patterns, and responses to boundary condition configurations.

### Mechanistic basis for parameter sensitivity differences

The LAM permeability *γ* acts as a vessel wall conductance parameter that controls the total volumetric exchange between the 1D vasculature and 3D tissue: at low *γ*, the vessel wall is a high-resistance barrier that limits fluid entry into the tissue, yielding net flow rates far below the perfusion-driven maximum. As *γ* increases, wall resistance becomes negligible and net flow asymptotically saturates at a level determined by vascular network architecture and boundary conditions rather than by wall resistance. This saturation behavior, which is observed consistently across all three boundary condition configurations (Cases 1, 2, and 3), confirms that at sufficiently high *γ*, the LAM transitions from a conductance-limited to a geometry-limited regime a result with direct implications for model identifiable. Once the model saturates, flow measurements show negligible sensitivity to *γ*.

In contrast, the SOI parameter *ε* modulates only the spatial extent over which a predefined terminal flow rate is distributed, without altering its total magnitude. Because the normalization condition (Eq. (5)) ensures that all flow exiting each terminal node is deposited in surrounding tissue regardless of *ε*, changes in sphere size affect how flow is spatially weighted but not how much of it enters the tissue globally. This explains the consistently gradual and non-saturating increase in SOI net flow with *ε* across all tested cases: the global flow reflects changes in the effective tissue conductance accessible to the terminal sources as the sphere radius expands, rather than a physical membrane resistance. This mechanistic distinction has practical implications: *γ* can in principle be inferred from total domain flow rate measurements alone, whereas *ε* is more directly constrained by regional perfusion measurements.

### Global flow behavior and boundary condition dependence

For the benchmark problem, the three boundary condition configurations revealed that global flow equivalence between LAM and SOI is configuration-dependent. The LAM saturation plateau persisted across all three configurations, confirming it is a formulation-level property independent of boundary conditions. The SOI flow increases almost linearly with the SOI radius. The Case 3 results (bilateral sinks at *y* = 0 and *y* = 40 mm) produced the largest net flow rates across all configurations, consistent with the greater combined drainage capacity of two active sink faces.

In Case 1 (sink at *x* = 0mm), both models asymptotically approach comparable net flow rates at high *γ* and large *ε*, suggesting that the global exchange capacity of the network governs both models similarly at their saturation limits. Moving the sink to *x* = 50 mm (Case 2) leaves both the LAM plateau and the SOI asymptotic flow rate close in magnitude to their Case 1 values, indicating that relocating the sink along the vessel axis has only a modest effect on the global saturation behavior of either model. This near-invariance suggests that the total exchange capacity of the coupled network, rather than the specific position of the drainage boundary relative to the vessel inlet, is the primary factor setting the asymptotic flow magnitude for both coupling formulations. In contrast, the dual-sink transverse configuration (Case 3) produces a markedly different regime. Both the LAM plateau and the SOI asymptotic value rise to roughly three times their Case 1 and Case 2 levels. Thus indicating that the orientation and number of drainage boundaries relative to the vessel network, is the dominant factor governing global saturation magnitude in both models; rather than sink location along a single axis. This pattern holds for both coupling mechanisms, showing that neither LAM’s distributed transmural exchange nor SOI’s terminal-node coupling is inherently insensitive to boundary geometry; rather, both respond similarly to changes in the overall drainage configuration while retaining their characteristic parametric dependence on*γ* and *ε*, respectively.

### The *γ*–*ε* equivalence as an implicit function

The equivalence relationship between *γ* and *ε* can be formalized as an implicit function defined by equal net flow rates. For a given vascular geometry and boundary configuration, define the equivalence locus as the set of parameter pairs (*γ, ε*) satisfying:

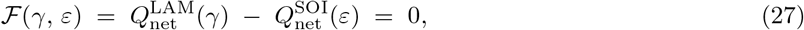

where 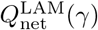 and 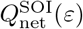 are the total domain flow rates produced by each model at the specified parameter values, with all other simulation inputs held fixed.

The global *Q*_net_ curves in Figs. 8.a, 10.a and 12.a provide the empirical basis for evaluating ℱ. In each panel, the LAM curve (blue) is a monotonically increasing, concave, saturating function of *γ*: it rises smoothly over roughly two decades of *γ* (from 10^−10^ up to ~ 10^−8^–10^−7^ m Pa^−1^s^−1^) and then plateaus, with no inflection point or S-shaped transition over the tested range *γ* ∈ [10^−10^, 10^−7^]. The SOI curve (orange), in contrast, increases close to linearly with *ε* across the tested range *ε* ∈ [5.8, 12] mm, without any sign of saturation.

Because both curves are monotonic over the tested ranges, Eq. (27) does not admit multiple *ε* solutions for a single *γ*: the equivalence locus is single-valued wherever it exists. What does change qualitatively across the parameter range is the *conditioning* of the equivalence, not its uniqueness. On the rising portion of the LAM curve (*γ* ≲ 10^−8^), a given change in *γ* produces a comparable, well-resolved change in *Q*_net_, so the corresponding equivalent *ε* is tightly constrained. Once *γ* enters the plateau (*γ* ≳ 10^−8^), however, further increases in *γ* produce almost no change in *Q*_net_: many distinct *γ* values in the plateau are consistent with essentially the same equivalent *ε*. In this regime *γ* is effectively unidentifiable from *Q*_net_ alone, even though the mapping itself remains formally single-valued.

Because the LAM plateau value differs across boundary configurations while the SOI curve’s dependence on *ε* is comparatively insensitive to sink placement, the *existence* of an equivalence locus is itself geometry-dependent. In Case 1 (sink co-located with the vessel inlet face, Fig. 8.a), the two curves converge to comparable *Q*_net_ magnitudes at their respective upper parameter limits, so an equivalence locus spans nearly the full tested range of both models. In Case 2 (sink on the opposite face, Fig. 10.a), the LAM plateau (~ 8.2 × 10^−10^ m^3^/s) is exceeded by the SOI curve well before *ε* reaches its upper tested limit (SOI grows from ~ 7.5 × 10^−10^ to ~ 1.5 × 10^−9^ m^3^/s over the same *ε* range), so a solution to Eq. (27) exists only over the lower portion of the tested *ε* interval. In Case 3 (bilateral sinks, Fig. 12.a), the SOI curve exceeds the LAM plateau (~ 2.8 × 10^−9^ m^3^/s) even at the *smallest* tested *ε* (SOI ranges from ~ 4 × 10^−9^ to ~ 8 × 10^−9^ m^3^/s) — here no exact global-flow equivalence exists anywhere in the tested parameter range, and matching *Q*_net_ would require extrapolating *ε* below the values used in this study.

Together, these observations show that *γ* and *ε* are not interchangeable parameters of equivalent physical models. They are fundamentally distinct coupling descriptors — *γ* characterizes a membrane conductance, *ε* characterizes a spatial distribution radius — and the range of *Q*_net_ values over which the two models can be matched at all is set by the boundary configuration, not by a fixed conversion factor. To the best of our knowledge, a fixed-ratio conversion between *γ* and *ε* is not achievable across the physiological parameter range relevant to tumor perfusion, drug delivery, or hemodynamic planning applications; any conversion established for one anatomical or boundary configuration should not be assumed to transfer to another.

### Regional perfusion and spatial heterogeneity in the benchmark model

In Case 1, both ROIs show clear but *opposite-signed* dependence on their respective coupling parameters (Fig. 8.b,c). The lower ROI’s outflow increases with *γ* (LAM: ~ 4.5 × 10^−11^ to a plateau of ~ 6.5 × 10^−11^ m^3^/s) and with *ε* (SOI: ~ 7 × 10^−11^ to ~ 1 × 10^−10^ m^3^/s), consistent with its role as a region close to the co-located sink: stronger coupling delivers more fluid that then drains locally. The upper ROI shows the reverse trend — outflow *decreases* as coupling strengthens (LAM: ~ 3.7 × 10^−11^ to ~ 1.5 × 10^−11^ m^3^/s; SOI: ~ 4.8 × 10^−11^ to ~ 1.1 × 10^−11^ m^3^/s), a 60–77% drop over the tested range. This reflects two compounding effects specific to Case 1’s geometry: more diffuse coupling deposits a growing share of the source outside the upper-ROI envelope, and because the sink sits on the same face that feeds the vessel network, stronger coupling increasingly short-circuits flow directly to the nearby sink rather than letting it reach the deeper terminal region. Regional perfusion in this configuration is therefore *not* insensitive to the coupling parameter — it is strongly and oppositely sensitive in the two ROIs, which is itself a practically important finding, since a single global-flow calibration would mask this local redistribution.

In Case 2 (Fig. 10), the lower ROI’s outflow remains positive throughout the tested range but is small and weakly decreasing with the coupling parameter (roughly 1–4 × 10^−12^ m^3^/s for both models) — this ROI sits transverse to the dominant vessel-to-sink axis and behaves as a near flow-through region rather than a net drain. The upper ROI, which now lies on the through-route between vessel inlet and the distal sink, shows the two models diverging in behavior: LAM outflow rises monotonically with *γ* toward a plateau (~ 1.3 × 10^−10^ to ~ 1.9 × 10^−10^ m^3^/s), while SOI outflow is non-monotonic, rising from ~ 2.5 × 10^−10^ m^3^/s at *ε* = 5.8 mm to a peak of ~ 2.9 × 10^−10^ m^3^/s near *ε* ≈ 7.5 mm before declining to ~ 2.5 × 10^−10^ m^3^/s at *ε* = 12 mm. The SOI peak marks the geometric crossover at which the sphere of influence first exceeds the distance from the terminal to the nearest ROI face; beyond that radius, an increasing fraction of each source sphere is deposited outside the ROI, lowering the net outflow measured through its bounding faces.

Under Case 3 (bilateral sinks, Fig. 12), the same qualitative asymmetry persists and is amplified. The LAM upper-ROI outflow rises with *γ* to a plateau near 4 × 10^−10^ m^3^/s — an order of magnitude larger than in Case 1 — reflecting the larger overall drainage capacity of the two-sink configuration. The SOI upper-ROI outflow is again non-monotonic, rising from ~ 1 × 10^−9^ m^3^/s at *ε* = 5.8 mm to a peak of ~ 1.1 × 10^−9^ m^3^/s near the same geometric crossover (*ε* ≈ 7.5 mm) before falling to ~ 7 × 10^−10^ m^3^/s at *ε* = 12 mm. That the SOI peak occurs at the same *ε* across Cases 2 and 3 supports a purely geometric interpretation — set by terminal-to-ROI-face distance, independent of sink placement — rather than a coupling-mechanism difference. The practical implication is that whenever the sink is well separated from the vessel inlets, LAM and SOI agree on the *direction* of the upper-ROI response to stronger coupling, but SOI delivers two-to-three times more flow through the region and does so non-monotonically, so the choice of coupling parameter carries substantially larger consequences for local perfusion predictions than for global ones.

### Regional perfusion analysis in the animal study

The swine liver animal study provided a physiologically complex validation context with a patient-specific hepatic arterial network reconstructed from CT hepatic arteriography. The regional analysis of model predictions (ROI 1 and ROI 2) revealed a substantial divergence in model predictions that extends qualitative benchmark-model findings to realistic clinical geometry.

For ROI 1, which is positioned adjacent to multiple vessel terminals, the SOI consistently predicts approximately 31–35% of total control-volume flow entering this region across the full range of arterial source efficiency *E* (7–21 voxels). Background Darcy perfusion—the passive flow driven by tissue pressure gradients—constitutes the dominant contribution (29–34%), while the direct arterial source accounts for only 0.3–3.3%, decreasing as *ϵ* increases because larger spheres distribute direct source contributions more broadly. This parametric stability reflects the SOI’s structural decoupling of total terminal flow rate from sphere size. In contrast, LAM predictions for the same region show dependence on *γ*, from approximately 10% at low *γ* to over 55% at high *γ*, driven by progressive recruitment of distributed wall exchange that disproportionately elevates perfusion estimates in tissue regions with close geometric proximity to vessel centerlines.

For ROI 2, positioned farther from vessel terminals, the SOI again predicts stable ≈ 14% total flow (perfusion ≈ 12–13%, source ≈1–3%) while LAM predictions range from ≈ 29% to ≈ 50% with increasing *γ*. Notably, at low *γ*, the LAM predicts more flow to ROI 2 than to ROI 1, inverting the spatial hierarchy that would be expected from proximity to vessel terminals. This inversion occurs because at low *γ*, wall exchange is globally suppressed and flow is dominated by the background pressure field established by the boundary conditions, which may direct flow through ROI 2 based on geometric connectivity rather than vascular proximity. As *γ* increases, vessel-proximal enhancement of ROI 1 progressively overcomes this geometric bias. The physiological implications for hepatic arterial chemoembolization (HACE) planning are clinically relevant: LAM predictions of drug delivery to a specific tumor region would exhibit sensitivity to vessel wall permeability, while SOI predictions would remain stable with corresponding differences in predicted treatment dose and efficacy.

The nondimensionalized *Q*_*net*_ comparison across inlet pressures of 80–100 mmHg corroborates these findings at the global scale: SOI predictions cluster in a narrow physiologically plausible band regardless of *E*, while LAM predictions span a markedly wider range depending on *γ*. Both models maintain physically consistent behavior within the reference flow range, confirming that both are valid representations of the hemodynamic physics but with different parameter dependencies.

### Clinical implications for interventional oncology

The hepatic arterial chemoembolization context of this animal study provides a clinically motivated framework for interpreting model selection guidelines. In HACE, the goal is to deliver embolic and chemotherapy agents through the hepatic arterial tree to achieve selective tumor necrosis while preserving surrounding parenchyma. Accurate prediction of regional perfusion heterogeneity is therefore essential for treatment planning, response prediction, and optimization of catheter placement. Our results suggest that the choice of coupling model fundamentally shapes these predictions in ways that cannot be resolved by global flow metrics alone.

The SOI model’s parametric stability offers a practical advantage for prospective treatment planning: regional perfusion estimates are robust to uncertainty in the sphere size parameter, and the model can be parameterized primarily from imaging-derived vascular geometry without requiring independent characterisation of vessel wall permeability. Conversely, the LAM’s strong *γ*-dependence provides richer mechanistic information when vessel wall permeability can be characterized—for example, from dynamic contrast-enhanced imaging or from prior knowledge of tumor vascular phenotype. In tumors with hyperpermeability from VEGF-driven angiogenesis [**?**], high *γ* values would dramatically redistribute flow toward vessel-proximal regions; in tumors with normalized vasculature following antiangiogenic treatment, low *γ* would produce more diffuse and geometrically controlled perfusion. The LAM is therefore a more mechanistically informative model when permeability characterization is available, while the SOI provides more reliable prediction uncertainty bounds when it is not.

Model validation in both frameworks should employ multiple regional perfusion measurements rather than global flow metrics alone, as the latter may fail to distinguish between models that predict very different spatial distributions of tissue perfusion. Multiparametric imaging approaches combining CT hepatic arteriography for vascular geometry with dynamic perfusion CT or MRI for tissue-level flow quantification would provide the regional ground-truth data needed to discriminate between model predictions.

Recent advances in imaging technology and computational power have made large-scale 3D-1D simulations increasingly feasible [**?, ?**], intensifying the need for systematic model comparison. Light-sheet fluorescence microscopy now enables whole-organ vascular network reconstruction with millions of resolved vessels [**?**], while clinical imaging modalities such as dynamic contrast-enhanced MRI provide spatially-resolved perfusion measurements suitable for model validation [**?**]. Simultaneously, advances in finite element methods, stabilization techniques, and preconditioning strategies have improved the computational efficiency and numerical stability of 3D-1D coupled solvers [**?, ?**]. These developments create an opportune moment to rigorously compare LAM and SOI model and establish quantitative guidance for model selection.

### Limitations

Several limitations of this study merit acknowledgment. First, the benchmark model employs a simplified vascular network with idealized geometry and boundary conditions; while this enables controlled comparison of the two coupling paradigms, it may not capture the full complexity of physiological vascular trees with thousands of branching vessels and multiscale hierarchy. Second, the absence of direct experimental validation of regional perfusion—via tracer kinetics, positron emission tomography, or ex vivo tissue measurements—means that absolute flow predictions of both models remain unvalidated against ground truth in the animal study. The lipiodol distribution observed post-treatment provides a qualitative spatial reference but does not quantitatively constrain model parameters or validate regional flow estimates. Third, the assumption of a single sink region for hepatic venous drainage in the animal study introduces uncertainty, as the true drainage geometry could not be resolved from the available CT imaging data; future studies with dedicated venographic imaging would reduce this uncertainty. Fourth, both models were implemented under the assumption of steady-state Darcy flow; time-dependent pulsatile effects and non-Newtonian blood rheology in small vessels were not considered. Finally, the benchmark model sensitivity analysis employed finite difference approximations, and the oscillatory behavior of LAM sensitivity at low *γ* may partially reflect numerical artifacts from the finite element discretization rather than pure model sensitivity.

## Conclusion

This study provides a systematic quantitative comparison of the Lateral Average Model and Sphere of Influence coupling paradigms in 3D-1D vascular-tissue simulations, with four principal conclusions. First, *γ* and *ε* play fundamentally different physical roles that preclude simple parameter interconversion. The LAM permeability *γ* controls vessel wall conductance and directly regulates total transmural exchange: 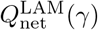 rises smoothly over roughly two decades of *γ* and then plateaus for *γ* ≳ 10^−8^, over the tested range *γ* ∈ [10^−10^, 10^−7^]. The SOI sphere radius *ε* instead controls only the spatial footprint of terminal flow delivery, producing a gradual, close-to-linear response with no saturation over the tested range *ε* ∈ [5.8, 12] mm. Because both responses are monotonic, matching *Q*_net_ between the two models does not require a multi-valued conversion, but it does require enough dynamic range in *ε* to reach the LAM plateau, and this is not always available: in Case 1 the two models converge to comparable *Q*_net_ at their respective upper parameter limits, in Case 2 the LAM plateau is undercut by the growing SOI curve within the tested *ε* range, and in Case 3 the SOI curve exceeds the LAM plateau even at its smallest tested *ε*, so no exact equivalence exists at all within the parameters used here. A fixed-ratio conversion between *γ* and *ε* is therefore not achievable across the physiological parameter range, and any conversion fitted to one boundary configuration should not be assumed to transfer to another.

Second, the two models differ in how identifiable their coupling parameter is from a flow measurement. Once LAM saturates (*γ* ≳ 10^−8^), *Q*_net_ becomes essentially insensitive to further increases in *γ*. This is visible directly in the animal-study comparison (Fig. 13), where the LAM mean 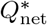 is nearly flat across four decades of *γ* at both tested inlet pressures, while its envelope is instead driven by the terminal-conductance parameter *γ*_*a*_. SOI, by contrast, shows a mild, non-saturating dependence of 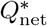 on *ε*^∗∗^ over the same range, with an envelope over (*γ*_*a*_, *γ*_*v*_) that is narrower than LAM’s and tightens further as *ε*^∗∗^ increases. In practical terms, *γ* becomes difficult to constrain from a global flow measurement once the LAM formulation saturates, whereas *ε* remains informative — and progressively better-constrained — across the full tested range. This makes SOI the more tractable formulation for parameter estimation from flow data when the LAM operating point is not known in advance to fall on the rising portion of its curve.

Third, regional perfusion predictions diverge substantially between models despite general agreement on which regions are net flow recipients. In the swine liver animal study, SOI consistently predicted ≈ 31–35% and ≈ 14% of total control-volume flow for ROI 1 and ROI 2 respectively, essentially independent of *ε*^∗∗^ (Figs. 14 and 15), while LAM predictions spanned ≈ 10–55% for ROI 1 and ≈ 29–50% for ROI 2 depending on *γ* — roughly a five-fold range for ROI 1. At low *γ*, LAM can even invert the expected perfusion hierarchy, predicting more flow to the more distant ROI 2 than to the vessel-proximal ROI 1, because wall exchange is globally suppressed and the background pressure field set by the boundary conditions dominates instead. The benchmark ROIs show the same qualitative pattern: Case 1 upper- and lower-ROI outflows change by 40–77% across the tested coupling range but in opposite directions, and the SOI upper-ROI response in Cases 2 and 3 is non-monotonic, peaking near *ε* ≈ 7.5 mm before declining. Both models therefore agree on coarse flow hierarchy but disagree substantially, and in model-specific ways, on the magnitude and even the parameter-dependence of regional perfusion.

**Fig 14.**
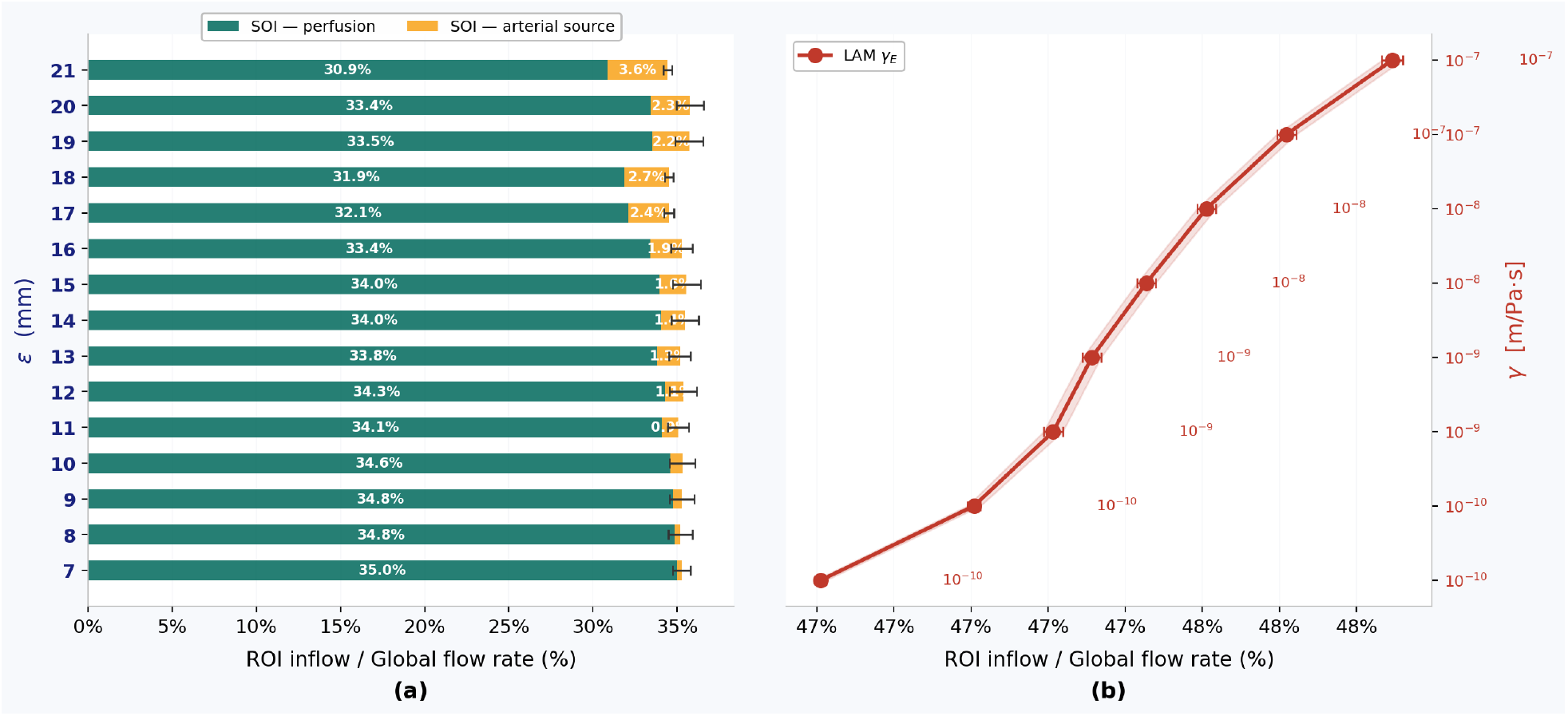
Breakdown of flow distribution in ROI 1 between the two models. The flow rate going in the ROI 1 is the percentage of total flow rate going in the CV. For the SOI model, the perfusion bar represents how much percentage of total CV flow rate is reaching ROI 1 based on neighboring tissue perfusion. The arterial source represents the percentage of total CV that is directly reaching the ROI 1 from the terminal vessels without ever flowing through any other tissue voxel. The percentage amount of CV flow reaching the ROI 1 in LAM is shown using the red plot and secondary y-axis shows the tissue-vessel permeability value.

**Fig 15.**
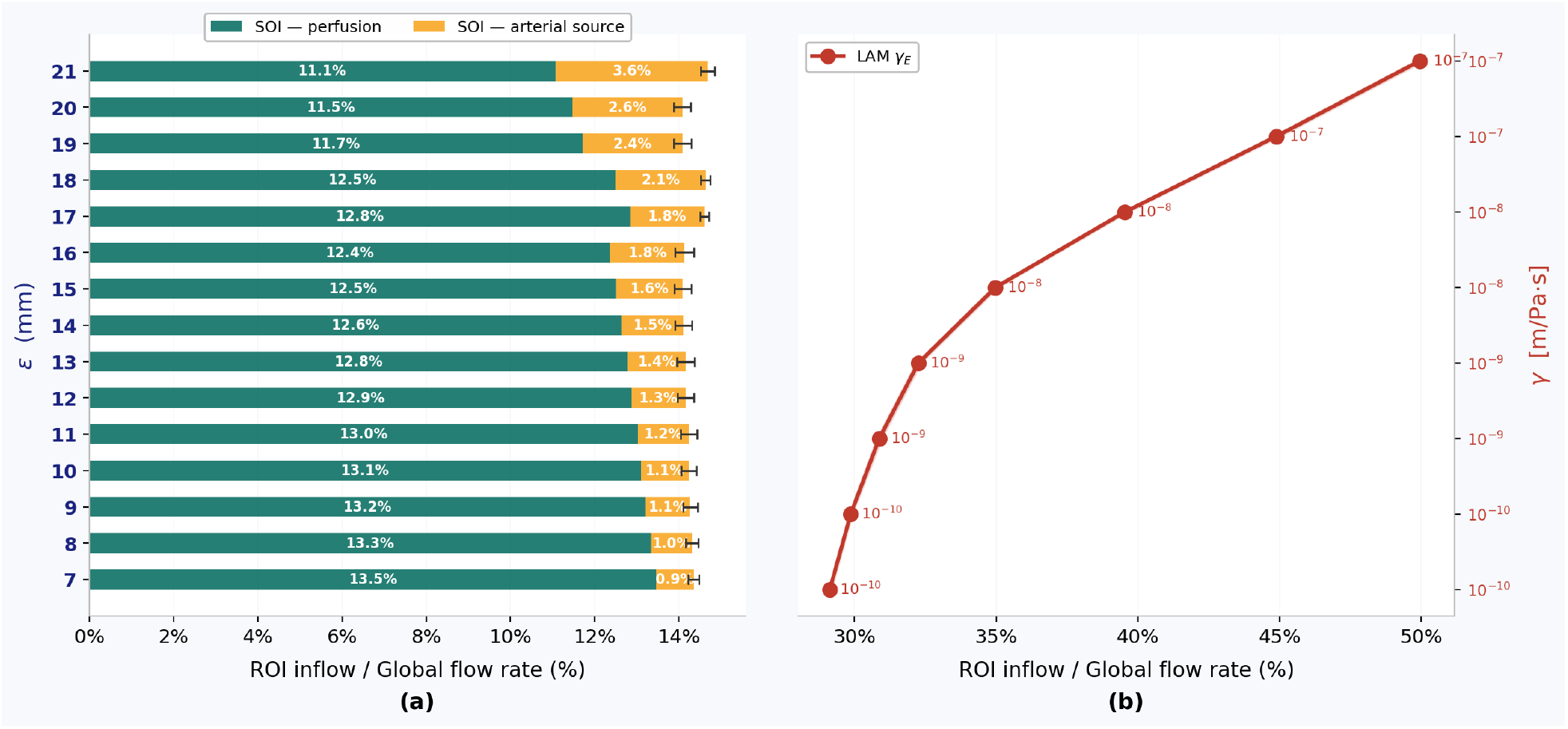
Breakdown of flow distribution in ROI 2 between the two models. The flow rate going in the ROI 2 is the percentage of total flow rate going in the CV. For the SOI model, the perfusion bar represents how much percentage of total CV flow rate is reaching ROI 2 based on neighboring tissue perfusion. The arterial source represents the percentage of total CV that is directly reaching the ROI 2 from the terminal vessels without ever flowing through any other tissue voxel. The percentage amount of CV flow reaching the ROI 2 in LAM is shown using the red plot and secondary y-axis shows the tissue-vessel permeability value.

**Fig 16.**
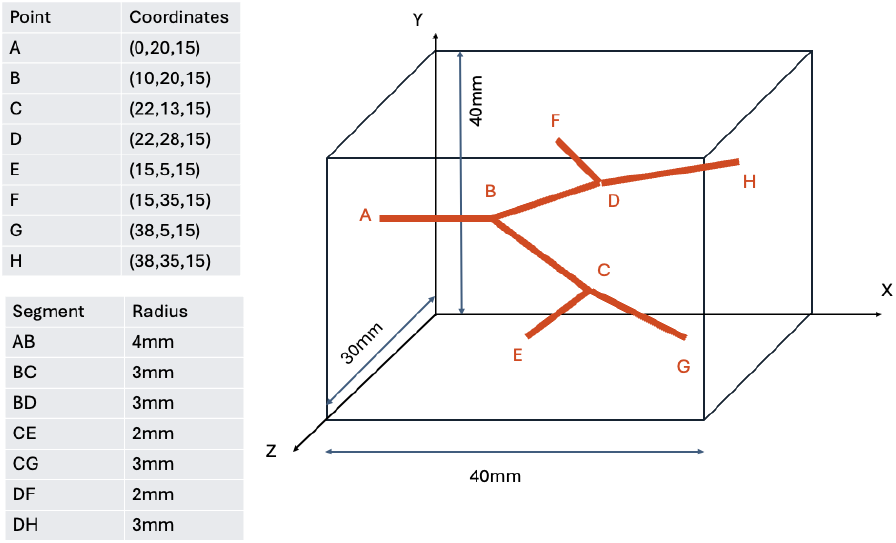
Coordinates of each graph node and radii of each vessel

Fourth, global flow equivalence between LAM and SOI is not a universal property but is specific to particular boundary-condition geometries. Relocating the sink from the vessel-inlet face (Case 1) to the opposite face (Case 2) leaves the LAM plateau essentially unchanged while the SOI curve grows further, widening the gap between the two models; adding a second, transverse sink pair (Case 3) raises both models’ flow rates roughly three-fold relative to Case 1 but does not restore an overlapping range between the LAM plateau and the tested SOI values. This shows that the two models encode different, boundary-geometry-dependent assumptions about the relationship between drainage configuration and vascular exchange, and that agreement (or disagreement) in global flow rate at one boundary configuration cannot be assumed to hold at another.

Together, these findings suggest application-specific guidelines: SOI is preferred for robust perfusion predictions when coupling parameters are uncertain, when sensitivity to parameters must be bounded for reliable optimization, or when regional flow stability is required for treatment planning. LAM is preferred when vessel wall permeability is a physiologically meaningful, identifiable, and measurable quantity particularly in inflammatory, oncological, or pharmacologically altered vascular beds where transmural permeability is the primary mechanism of interest. In all cases, model validation should employ multiple regional perfusion measurements rather than global flow metrics alone, as global equivalence can mask large regional discrepancies with direct consequences for predicted drug delivery, oxygenation, and therapeutic response.

## Supporting information

**S1 Fig. Coordinates of each graph node and radii of each vessel**.

## Acknowledgments

The authors acknowledge the support of NIH grant 1R01CA301555-01. Riviere acknowledges support from NSF DMS 2513092 and NSF DMS 2231482.

## Notes

### Competing Interest Statement

The authors have declared no competing interest.

